# Predicting the First Steps of Evolution in Randomly Assembled Communities

**DOI:** 10.1101/2023.12.15.571925

**Authors:** John McEnany, Benjamin H. Good

**Author notes:** Correspondence should be addressed to: B.H.G.

## Abstract

Microbial communities can self-assemble into highly diverse states with predictable statistical properties. However, these initial states can be disrupted by rapid evolution of the resident strains. When a new mutation arises, it competes for resources with its parent strain and with the other species in the community. This interplay between ecology and evolution is difficult to capture with existing community assembly theory. Here, we introduce a mathematical framework for predicting the first steps of evolution in large randomly assembled communities that compete for substitutable resources. We show how the fitness effects of new mutations and the probability that they coexist with their parent depends on the size of the community, the saturation of its niches, and the metabolic overlap between its members. We find that successful mutations are often able to coexist with their parent strains, even in saturated communities with low niche availability. At the same time, these invading mutants often cause extinctions of metabolically distant species. Our results suggest that even small amounts of evolution can produce distinct genetic signatures in natural microbial communities.

## Main Text

Microbes often live in ecologically complex communities containing hundreds of coexisting species (1–4). As residents of these communities compete with each other, they can evolve over time by acquiring mutations (5–7). These evolutionary changes can alter the ecological interactions between species, driving shifts in community composition (8–10). Conversely, the community also creates the context in which organisms evolve, by influencing the structure of the adaptive landscape (11–15). Understanding how the community influences these evolutionary paths (and vice versa) is a crucial step toward predicting and controlling microbial ecosystems.

Longstanding conceptual models suggest that community structure can impact evolution in different ways. Some models predict that the rate of evolution should decline in larger communities, as more niches are filled by other species (7, 16–18). Others have proposed that diverse communities could create new opportunities for local adaptation, by sup-pressing key competitors or creating new niches through cross-feeding (11–14, 19, 20). These conceptual models also make different predictions about how adaptive mutations will impact their community when they invade. Some mutations will replace their parent strain, while others can encroach on other species (9, 21) or diversify into coexisting lineages (5, 6, 22). Each of these behaviors has been observed empirically, yet it remains challenging to predict which should dominate in a given community.

The source of this challenge lies in the niches inhabited by different species, and how mutations alter or move between them. While much progress has been made in small communities where the relevant niches can be explicitly defined (21–24), it is difficult to extend this approach to larger communities like the gut microbiome, where species can compete for many different combinations of resources. In this high-diversity limit, even basic questions about the effects of community structure remain unresolved: how does the benefit of a mutation depend on the diversity of the community and the metabolic overlap between its members? Do mutations primarily compete with their parent strain, or do they continue to stably diversify, as suggested by recent evidence from the gut microbiome (5, 6, 25) and some *in vitro* communities (26, 27)?

Resource competition models provide a mechanistic framework to address these questions (28–32). In these models, niches are not defined in advance, but emerge organically through differences in resource consumption (33). An extensive body of work has used this framework to investigate the process of community assembly, where species compete to colonize a new environment (28, 34–37). These model communities can recapitulate some large-scale features of natural (38–40) and experimental ecosystems (39–42). By contrast, the evolutionary dynamics that emerge from resource competition are difficult to model with traditional community assembly theory. While some studies have started to explore these effects, previous work has mostly focused on small communities (30, 31, 43) or the long-term states attained over infinite evolutionary time (30, 44). Both approaches are poorly suited for understanding how a focal species evolves in different community backgrounds, which is the scenario most accessible in experiments. To address this gap, we develop a theoretical framework for predicting the initial steps of evolution in an assembled community with many coexisting species. By extending random matrix approaches from community assembly theory, we derive analytical predictions that describe how the fitness benefits and fates of new mutations scale with the diversity and metabolic overlap of the surrounding community, enabling quantitative tests of the conceptual models above.

### Modeling first-step mutations in randomly assembled communities

To study the interplay between ecological competition and new mutations in a mathematically tractable setting, we turn to a simple resource competition model (28–32, 36, 45) where microbes compete for ℛ ≫ 1 substitutable resources that are continuously supplied by the environment (Fig. 1A). Each strain *µ* in the community is characterized by a resource utilization vector 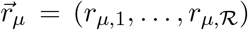, which describes how well it can grow on each of the supplied resources. While our model allows for simple forms of cross-feeding (see below), we neglect additional factors like regulation (46), resource sequestration (47), or antagonistic interactions (48), allowing us to focus on the basic evolutionary pressures imposed by resource competition alone.

**Figure 1:**
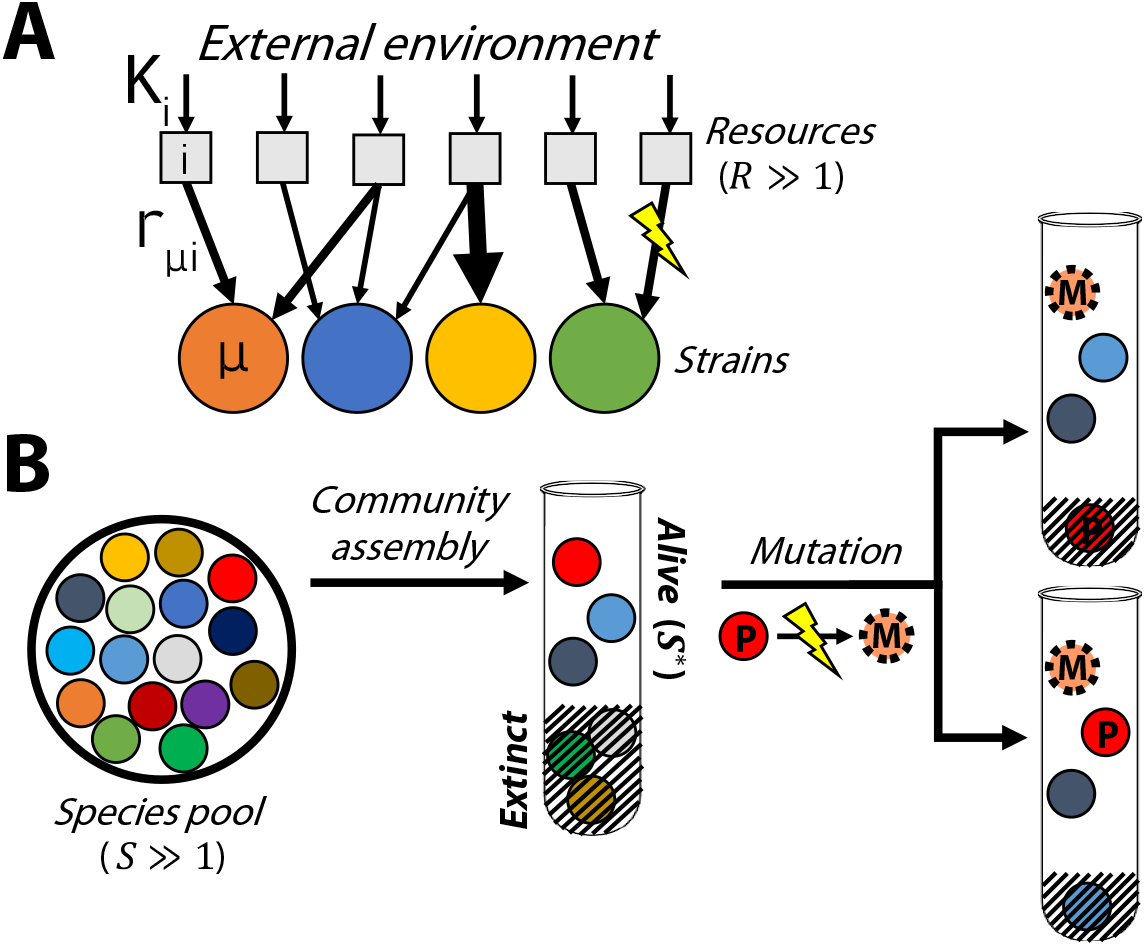
Modeling the first steps of evolution in a randomly assembled community that competes for substitutable resources. **(A)** Microbial strains compete for ℛ resources that are continuously supplied by the environment at rates *K*_*i*_. Each strain *µ* has a characteristic set of uptake rates *r*_*µ,i*_ (arrows), which can be altered by further mutations. **(B)** A local pool of 𝒮 initial species, whose phenotypes are randomly drawn from a common statistical distribution, self-assembles into an ecological equilibrium with 𝒮^*^ ≤ 𝒮 surviving species (left). A new mutation (*M*) arises in one of the surviving species (*P*); if the mutation provides a fitness benefit, its descendants can either replace the parent strain (top right) or stably coexist with the parent, potentially driving another species to extinction (bottom right).

In the simplest version of our model, the concentrations of the microbial strains 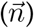 and abiotic resources 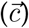 can be described by the coupled system of equations,

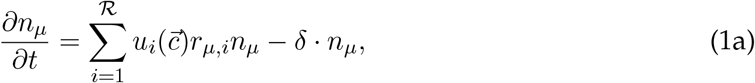

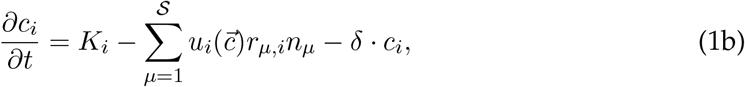

where *δ* is the dilution rate, *K*_*i*_ is the external supply rate of resource *i*, and 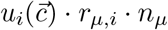 is the total uptake rate of resource *i* by species *µ* (SI Section 1.1). When the dilution rate is sufficiently low, or biomass sufficiently high, we can coarse-grain over time to obtain an effective model for the relative abundances of the strains (*f*_*µ*_),

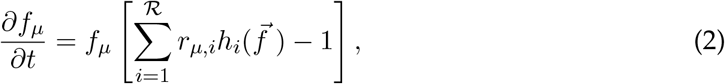

where time is now measured in generations and 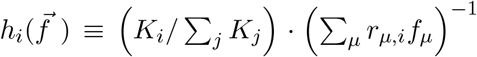 denotes the local availability of resource *i* (Fig. S1; SI Section 1.2). Simple forms of cross-feeding (39, 49) can also be included in this model by replacing the external supply rates 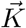 with an effective value that accounts for internal metabolic conversion (SI Section 1.3).

Previous work has used this model to study the process of community assembly (28, 36, 37), where 𝒮 initial strains arrive in a new environment and form an ecologically stable community containing 𝒮^*^ ≤ ℛ survivors (Fig. 1B, left). The steady-state values of 𝒮^*^ and 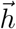 depend on the environmental supply rates 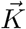 and the resource preferences 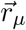 of the initial strains. While it is difficult to measure these kinetic parameters directly, past research has shown that emergent features of large ecosystems can often be mimicked by randomly drawing the uptake rates of each strain from a common statistical distribution (28, 29, 34–36).

For concreteness, we will initially focus on the binary resource usage model from Ref. (36), in which each strain has an independent probability ℛ_0_ /ℛ of utilizing each resource, and an overall uptake budget *X*_*µ*_ ≡log ∑_*i*_ *r*_*µ,i*_ that is independently drawn from a Gaussian distribution (SI Section 1.4). However, our results will apply for a variety of different sampling schemes, which we verify in Figures S3, S8, and S9 and SI Section 4.

Individual realizations of this model produce assembled communities with similar numbers of surviving species, which can be predicted using methods from the physics of disordered systems (28, 29, 36; Fig. 2B; SI Section 3). In our analysis below, it will be convenient to treat the expected number of surviving species as an input parameter, and classify the assembled communities as a function of their niche saturation, 𝒮^*^/ ℛ (SI Section 3.2). This re-parameterization allows for easier generalization to other community assembly models, since communities with similar levels of niche saturation will often have similar statistical properties, even if they resulted from different sampling distributions (29, 35, 36).

**Figure 2:**
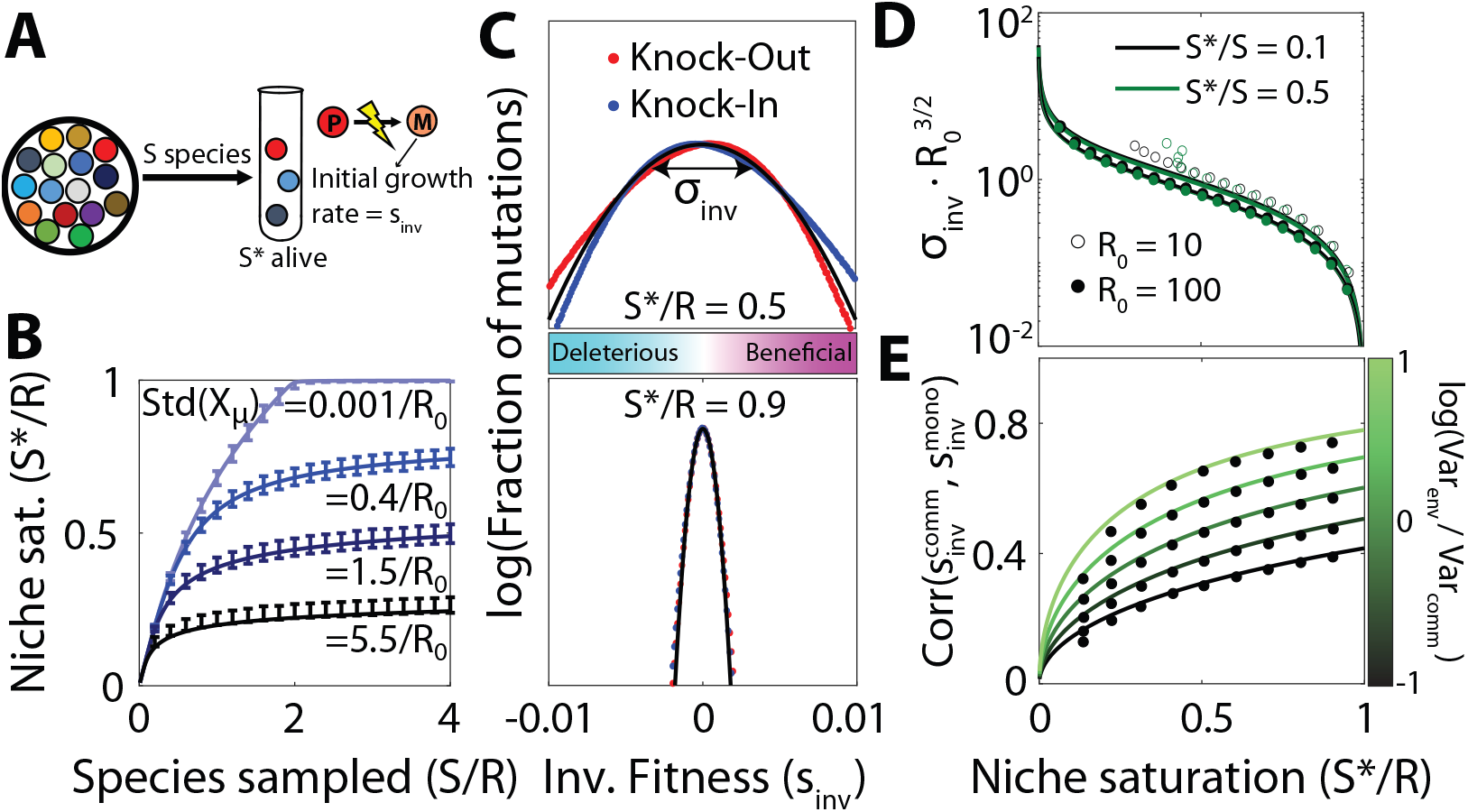
Distribution of fitness effects of first-step mutations as a function of community complexity. **(A)** As in Fig. 1, a community is assembled from 𝒮 initial species, leaving 𝒮^*^ alive at equilibrium. The surviving species produce mutations, whose invasion fitness *s*_inv_ is equal to their initial relative growth rate. **(B)** Number of surviving species 𝒮^*^ as a function of the sampling depth 𝒮 and the standard deviation of the total uptake budget among sampled species, Std(*X*_*µ*_). Curves show theory predictions from SI Section 3.1, while points show means and standard deviations over 10^3^ simulation runs with ℛ = 200, ℛ_0_ = 40, and uniform resource supply. **(C)** Distribution of fitness effects of knock-out (and knock-in) mutations with Δ*X* = 0 in communities with two different levels of niche saturation (𝒮^*^/ℛ). Black curve shows the theoretical predictions from Eq. (5), while dots represent a histogram over all possible strategy mutations in 10^3^ simulation runs using the same parameters as panel B, with 𝒮^*^/𝒮 = 0.1. **(D)** The width *σ*_inv_ of the distribution of fitness effects in panel C as a function of niche saturation, for various values of sampling permissivity 𝒮^*^/𝒮 and per-species resource usage ℛ_0_. Curves show the theoretical predictions from Eq. (5), while the dots show the average over 10^3^ simulation runs. **(E)** Pearson correlation between the fitness effect of a mutation in the community and its fitness effect in monoculture, for different values of niche saturation and scaled variation in resource supply rates, 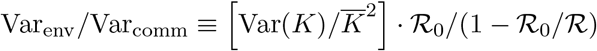. Curves show the theoretical predictions from SI Section 4.1, while points show the average over all mutations in 10^3^ simulated communities with parameters the same as panel C.

To account for evolution, we model the first mutational steps that occur in a randomly assembled community (Fig. 1B). This scenario might apply to the initial phases of *in vitro* passaging experiments (12, 26), or recently colonized gut microbiomes (50, 51). Through much of our analysis, we will focus on a particularly simple class of “knock-out” mutations, where the mutant loses its ability to consume one of its resources (*r*_*i*_ → 0). We assume that the cell can compensate for this deletion by increasing the uptake of other resources in its repertoire, but this compensation may not be perfect, corresponding to a shift in the effective uptake budget (*X* → *X* + Δ*X*). We also consider “knock-in” mutations, where a strain gains the ability to consume a resource (e.g. through horizontal gene transfer; 25), as well as more general changes that influence the uptake rates of multiple resources simultaneously 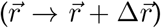. Following previous work (30, 32), it will be convenient to classify these mutations by their impact on the strain’s total resource uptake rate *X* ≡log ∑_*i*_ *r*_*i*_ and its normalized resource consumption strategy 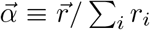, which describes the relative effort devoted to importing each of the resources. A general mutation will therefore involve a shift in one or both of these parameters (*X* → *X* + Δ*X* and 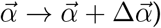.

While the phenotypes of these mutants are simple, their fitness consequences will depend on the local environment, which is shaped by the other resident strains. Using the assembled communities as a backdrop allows us to quantify how this evolutionary landscape varies with the size and composition of the surrounding community.

### Distribution of fitness effects of mutations in newly assembled communities

The rate of evolution in a newly assembled community will depend on the supply of beneficial mutations. This landscape is often summarized by the local distribution of fitness effects (DFE), denoted by *ρ*_*µ*_(*s*), which gives the relative probability that a mutation in focal strain *µ* will have an invasion fitness *s*. The shape and scale of *ρ*_*µ*_(*s*) determine the availability of beneficial mutations, and therefore the rate of evolutionary change. Our resource competition framework allows us to ask how these features depend on the composition of the larger community.

For a community at ecological equilibrium, a mutation that arises in a resident strain *µ* and changes its resource uptake phenotype to a new value *X*_*µ*_ + Δ*X* and 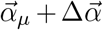 will have an invasion fitness,

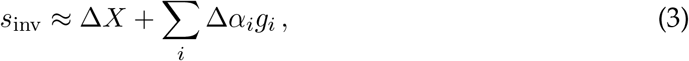

where 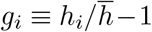 is the excess availability of resource relative to the ecosystem average, 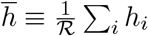 (30) (SI Section 1.5). Equation (3) shows that the invasion fitness of a mutation that only affects the overall uptake budget of a strain 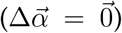 is independent of the surrounding community. In contrast, the benefits of mutations that change the resource consumption strategy of a strain will depend on the interactions between species, which are mediated by the values of the resource availabilities, *g*_*i*_. For example, if a resource has a lower relative availability (*g*_*i*_ *<* 0), then it is not worth devoting a portion of its uptake budget to consume it, and a knock-out mutation for that resource should be beneficial.

Replica-theoretic calculations similar to Ref. (36) allow us to predict the distribution of the *g*_*i*_ for a typical assembled community when ℛ ≳ ℛ_0_ ≫ 1 (SI Section 3.2). In large communities, the excess resource availabilities are well-approximated by a collection of Gaussian random variables,

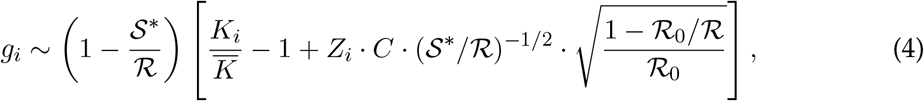

where 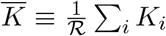 is the average resource supply rate, *C* is an 𝒪(1) factor that depends on the sampling depth 𝒮/𝒮^*^, and the *Z*_*i*_ are uncorrelated standard Gaussians. Analogous expressions for other sampling distributions can be obtained by replacing the ℛ_0_-dependent factor with the corresponding spread in 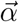 (e.g. SI Section 4.4). This mean-field solution requires that the resources are supplied at comparable rates 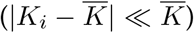 and each resource is utilized by a substantial number of surviving strains (𝒮_0_ 𝒮^*^≫ 𝒮 SI Section 3.5; Fig. S3D). When these conditions are satisfied, we can use Eqs. (3) and (4) to calculate the distribution of fitness effects *ρ*_*µ*_(*s*) by aggregating over mutations with different values of 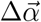 and Δ*X*.

To understand how the community influences the DFE of a focal species, it is helpful to begin by considering the simplest case, where the resource supply is uniform 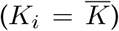 and mutations have no direct costs or benefits (Δ*X* = 0). In this case, Eqs. (3) and (4) imply that *ρ*_µ_(*s*) will also follow a Gaussian distribution with mean zero and standard deviation

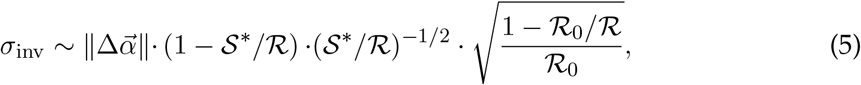

where 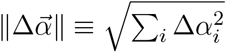 is the magnitude of the phenotypic change produced by the mutation. Examples for knock-out and knock-in mutations (where 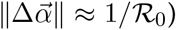 are shown in Fig. 2C-D. Since Eq. (5) does not explicitly depend on the parent strain *µ*, it implies that the DFEs should be statistically similar for all strains in the community (Fig. S2). Furthermore, since the details of the mutations only enter through their overall magnitude 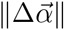, this implies that knock-out and knock-in mutations — as well as multi-resource mutations with the same value of 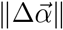 — will have statistically similar DFEs (Fig. 2C).

Equation (5) shows that the community influences the DFE of a focal strain primarily through the degree of niche saturation, 𝒮^*^/ℛ. The magnitude of the typical fitness effect approaches zero as 𝒮^*^/ℛ increases (Fig. 2D), which is consistent with the idea that the rate of evolution will be slower in communities where more niches are already filled, since the establishment probability of a beneficial mutant is proportional to its fitness effect (52).

However, since the mean of the DFE is still centered at *s* = 0, the overall fraction of beneficial mutations remains constant as the niche saturation increases (Fig. 2C), even though a smaller number are likely to survive genetic drift. This implies that surviving organisms are not necessarily at a local evolutionary optimum, where any change to their resource consumption strategy tends to be deleterious.

However, this symmetry between the frequency of beneficial and deleterious mutations critically depends on the assumption of perfect trade-offs (Δ*X* = 0), which will not always hold in practice. For example, a beneficial mutation could halt the production of an enzyme which is used for metabolizing a low-availability resource, while leaving the expression of other enzymes in that now-defunct pathway intact – resulting in a net cost to pure fitness. If all mutations carried such a direct cost (Δ*X* < 0), then Eq. (3) implies that the entire DFE would shift to the left by a constant amount −|Δ*X*|. If this shift is larger than the typical spread of the DFE (|Δ*X*| ≫*σ*_inv_), then the beneficial tail of *ρ*_*µ*_(*s*) will approach an exponential shape, whose height and width will both strongly decline with the degree of niche saturation 𝒮^*^/*ℛ*. Thus, the availability of beneficial mutations can sensitively depend on the genetic architecture of the strain’s resource consumption rates.

Similar results apply when the resources are supplied at different rates (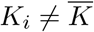; SI Section 4.1; Fig. S7). In this case, the fitness effect of a mutation will depend on both the community context and the external environment, as encoded by the resource availabilities in Eq. (4). The overall magnitude of the environmental contribution declines as niche saturation 𝒮^*^/*ℛ* increases, consistent with previous work showing that larger communities self-organize to “shield” their member species from the external environment (30, 32, 36). Interestingly, however, Eq. (4) shows that the *relative* impact of 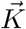 on the fitness effect a mutation actually increases with the degree of niche saturation, so the external environment can still exert an influence on the overall direction of the fitness landscape. This effect is strikingly illustrated when we compare the fitness effect of each mutation with its expected value in the absence of the community (Fig. 2E). We find that the correlations between the t<u>wo DFEs c</u>an be substantial when the coefficient of variation in *K*_*i*_ exceeds a critical value, 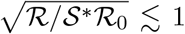. This indicates that environmental shielding is often incomplete: the net direction of natural selection can be preserved across community backgrounds, even when inter-specific competition exerts a strong effect on the local resource availabilities.

### Ecological diversification in large communities

While the invasion fitness of a mutation describes its initial growth rate, a successful variant will eventually reach a size where it starts to impact the other members of the community. Some of these mutants will eventually replace their parent strain, due to the principle of competitive exclusion (28). Others can stably coexist with their parents by exploiting a different ecological niche (30). How does the frequency of these *in situ* ecological diversification events depend on the composition of the surrounding community?

We can analyze the probability of mutant-parent coexistence in our model by recasting it as the outcome of two correlated assembly processes. First, an initial ecosystem *E*_0_ is formed through our standard community assembly process (Fig. 1B, left). Then, a second ecosystem is formed when one of the surviving species in *E*_0_ produces a beneficial mutant *M*, and the combined community *E*_0_ + *M* is allowed to reach its new ecological equilibrium (Fig. 1B, right). The latter event requires that the parent strain has a positive relative abundance in the first ecosystem 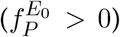, and that the mutant survives in the second ecosystem 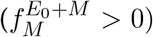. The probability that the mutant coexists with its parent can then be expressed as a conditional probability,

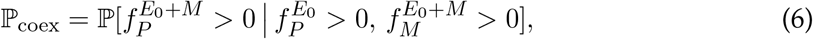

which averages over the random resource uptake rates in the initial community, as well as the random effect of the adaptive mutation.

The correlated assembly process in Eq. (6) is challenging to analyze with existing methods (37), since the mutant and parent phenotypes are closely related to each other. Fortunately, we will show that we can often approximate the coexistence probability by considering a third community assembly process, in which the mutant and parent are introduced simultaneously with the other species. This “simultaneous assembly” approximation differs from the two-ecosystem model in Eq. (6), since the final community can contain “rescued” species that would not have survived in *E*_0_ before the mutant strain invaded. However, simulations indicate that these differences result in only small corrections to the coexistence probability across a broad range of parameters (SI Section 2.1, Fig. S4), so this is often a reasonable approximation.

When the simultaneous assembly approximation holds, we can evaluate the coexistence probability by extending the replica-theoretic calculations used for Eq. (4) (SI Section 3.3). We find that coexistence can be expressed as the probability that the invasion fitness of the mutant lies below a critical fitness threshold,

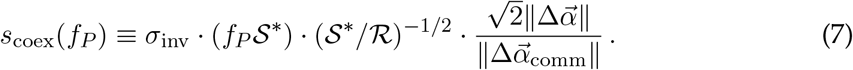

where *f*_*P*_ is the relative abundance of the parent strain, 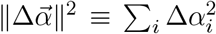 is the pheno-typic effect size of the mutation, and 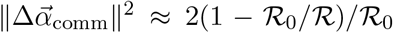 is the equivalent spread between random pairs of strains in the community. Averaging over the abundance of the parent strain (SI Section 3.4) yields a corresponding expression for the coexistence probability as an integral over the DFE in Fig. 2C,

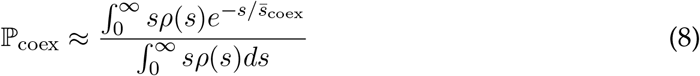

where 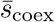 is the coexistence threshold for a typical genetic background (*f*_*P*_ *∼* 1/ 𝒮^*^). For mutations with modest phenotypic effects, such as a single resource knock-out 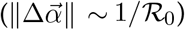, the magnitude of 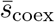 will be much smaller than the typical spread in the DFE in Fig. 2C. This implies that only mutations with anomalously low invasion fitnesses will have an appreciable chance of coexisting with their parent (Fig. 3A).

**Figure 3:**
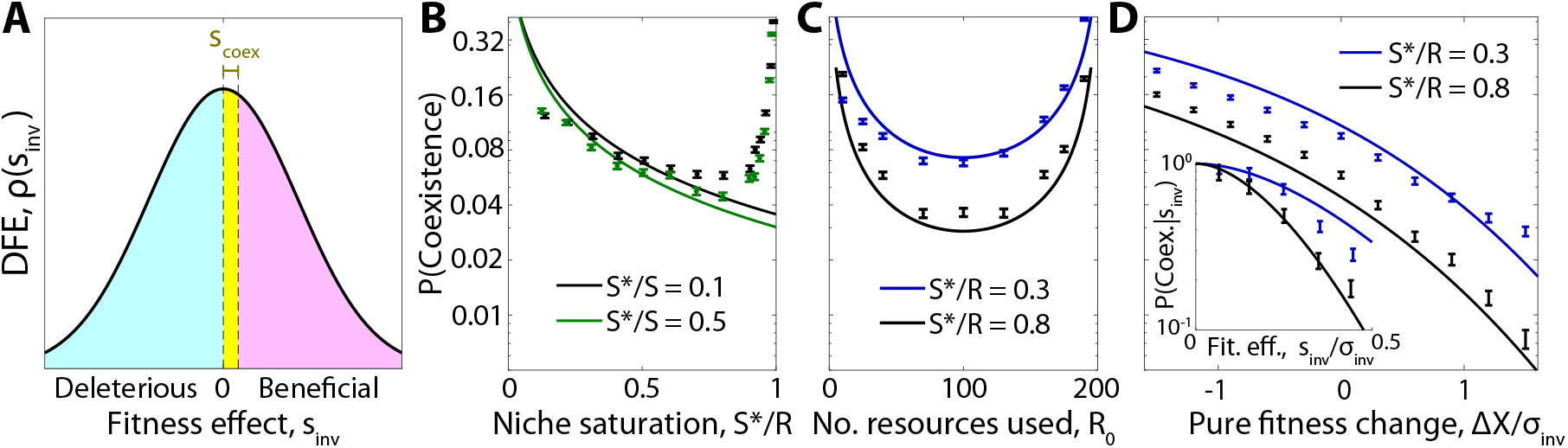
Ecological diversification in large communities. **(A)** Schematic showing the subset of mutations that are able to stably coexist with their parent strain. Coexistence occurs in the yellow region where the invasion fitness is below a critical threshold *s*_coex_, which depends on the abundance of the parent and the phenotypic effect size of the mutation. **(B-D)** Probability that a successful knock-out mutant coexists with its parent strain as a function of (B) the niche saturation 𝒮^*^/ℛ, (C) the typical number of resources used per species ℛ_0_, and (D) the change in overall uptake budget of the mutant Δ*X*. Inset shows the dependence on the total invasion fitness *s*_inv_. In all panels, curves show the theory predictions from SI Sections 3.3 and 3.4, while points show means and standard errors over 10^4^ simulation runs with base parameters ℛ= 200, ℛ_0_ = 40, 𝒮^*^/𝒮 = 0.1, and uniform resource supply.

For a mutation with perfect trade-offs (Δ*X* = 0), the integral in Eq. (8) reduces to a simple form,

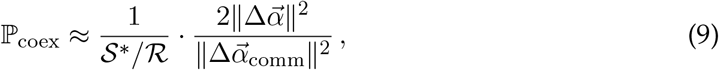

which depends on the niche saturation 𝒮^*^/ℛ and the phenotypic effect size of the new mutation.

Equation (9) shows that the coexistence probability is largest for small values of 𝒮^*^/ℛ (Fig. 3A), consistent with the idea that diversification is easier when there are many open niches left to exploit, as in an adaptive radiation (20). Interestingly, however, we find that the coexistence probability does not vanish as the community approaches saturation (𝒮^*^/ℛ → 1), but instead plateaus at a nonzero value. This is true even in fully saturated communities (𝒮^*^ = ℛ), where other species must be driven to extinction when the successful mutant invades.

In this regime, the coexistence probability is most strongly determined by the phenotypic effect size of the mutation. A mutation that changes the resource consumption strategy by an infinitesimal amount 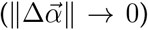 can never coexist with its parent, while a mutation with 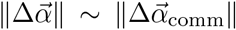 has a much higher probability of coexistence. Single-resource knockouts fall between these two extremes, with the coexistence probability scaling inversely with ℛ_0_ (the average number of resources utilized per strain), rather than the total number of resources consumed by the community. This suggests that *in situ* ecological diversification can continue to occur even in large communities containing many species and resources (Fig. 3B and Fig. S5).

We see that simulation results generally support the theoretical predictions in Eq. (9), but start to exceed this value for communities very close to full saturation. The source of this deviation coincides with the breakdown of our “simultaneous assembly” approximation above, which allowed the invading mutant to “rescue” some species that had previously gone extinct. The deviations disappear if extinct species are allowed to re-invade (Fig. S4), suggesting that competition from rescued species plays a significant role in our theory for extremely high levels of niche saturation. This rescuing effect could be relevant in some natural settings, e.g. if the species pool is maintained in a separate spatial niche with frequent migration back into the community. Regardless, our results in Eq. (9) provide a lower bound on the coexistence probability in large ecosystems, suggesting that diversification could be even more common under some conditions.

In reality, most mutations that change an organism’s resource consumption strategy are unlikely to be perfect trade-offs. More generally, we find that mutations that carry a direct cost or benefit Δ*X* change the coexistence probability in Eq. (9) by the factor

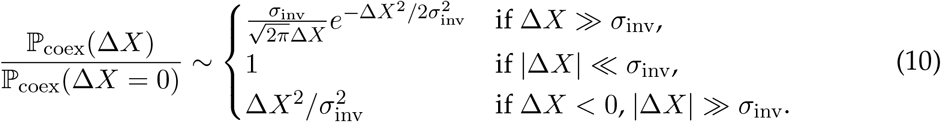

which shows that direct costs or benefits begin to exert an effect when Δ*X* becomes comparable to *σ*_inv_. However, the direction of this effect is somewhat counterintuitive: strategy mutations that carry a direct fitness cost (Δ*X <* 0) are more likely to coexist with their parent, provided that their overall invasion fitness is still positive. Conversely, mutations with strong direct benefits (Δ*X >* 0) are more likely to drive their parent strain to extinction (Fig. 3B). These differences arise because nonzero values of Δ*X* change the typical invasion fitnesses of successful mutants. The coexistence probability in Eq. (8) is highly sensitive to these shifts, as mutants with an invasion fitness below 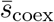 are much more likely to coexist with their parent (Fig. 3A,D). This is a reflection of the fact that diversification is driven by a narrow range of mutants with fitness high enough to establish but small enough to coexist.

We can gain some intuition for this effect by considering a loss-of-function mutation for a resource that is currently overutilized by the population (*g*_*i*_ *<* 0). Equation (2) shows that the growth rate of this variant (relative to its ancestor) is given by ∼ *g*_*i*_(*t*). As the mutant grows in abundance, it begins to replace its parent, which tends to reduce the overall utilization of the target resource (*∂*_*t*_*g*_*i*_ *>* 0). If *g*_*i*_ reaches zero before the parent strain goes extinct, then the mutant will coexist with its parent, having lost its initial growth advantage. This coexistence is most likely to happen if the parent strain has high abundance, or if |*g*_*i*_| (and therefore *s*_inv_) is initially small (Fig. S6).

The same argument applies for more complex mutations which affect multiple resources (Fig. S5). It also explains why mutations with small phenotypic effects (e.g. a pure fitness mutation with no strategy change) cannot coexist with their parent. If the mutant and parent have near-identical resource consumption strategies, the mutant’s invasion produces very little negative feedback on its growth rate relative to its parent, making it likely the parent strain will be driven to extinction (Fig. S5B). Similar logic applies for extensions of our basic model, including non-uniform resource supply rates and alternative community assembly schemes (Figures S3, S8, and S9; SI Section 4). Together, these results suggest that *in situ* diversification could be common even in large and saturated communities, particularly for mutants on abundant backgrounds with lower-than-expected invasion fitnesses.

### Successful mutations drive extinctions in other niches

In addition to displacing their parents, successful mutants can also drive other species in the community to extinction. This is particularly evident in saturated communities (𝒮^*^ = ℛ), where competitive exclusion implies that the invasion of a new strain must be accompanied by extinction of at least one other. Figure 4B shows that the number of extinctions steadily rises with the degree of niche saturation, exceeding 1% of the initial community (∼ 1-10 species) for many combinations of parameters. Moreover, these extinctions are not completely independent of each other, since they are somewhat overdispersed compared to a simple Poisson expectation (Fig. 4B, inset). Previous work has suggested that even distantly related strains could exhibit correlated dynamics if their resource consumption strategies are anomalously similar to each other (26). Could such hidden metabolic similarity be responsible for driving the extinction events above?

**Figure 4:**
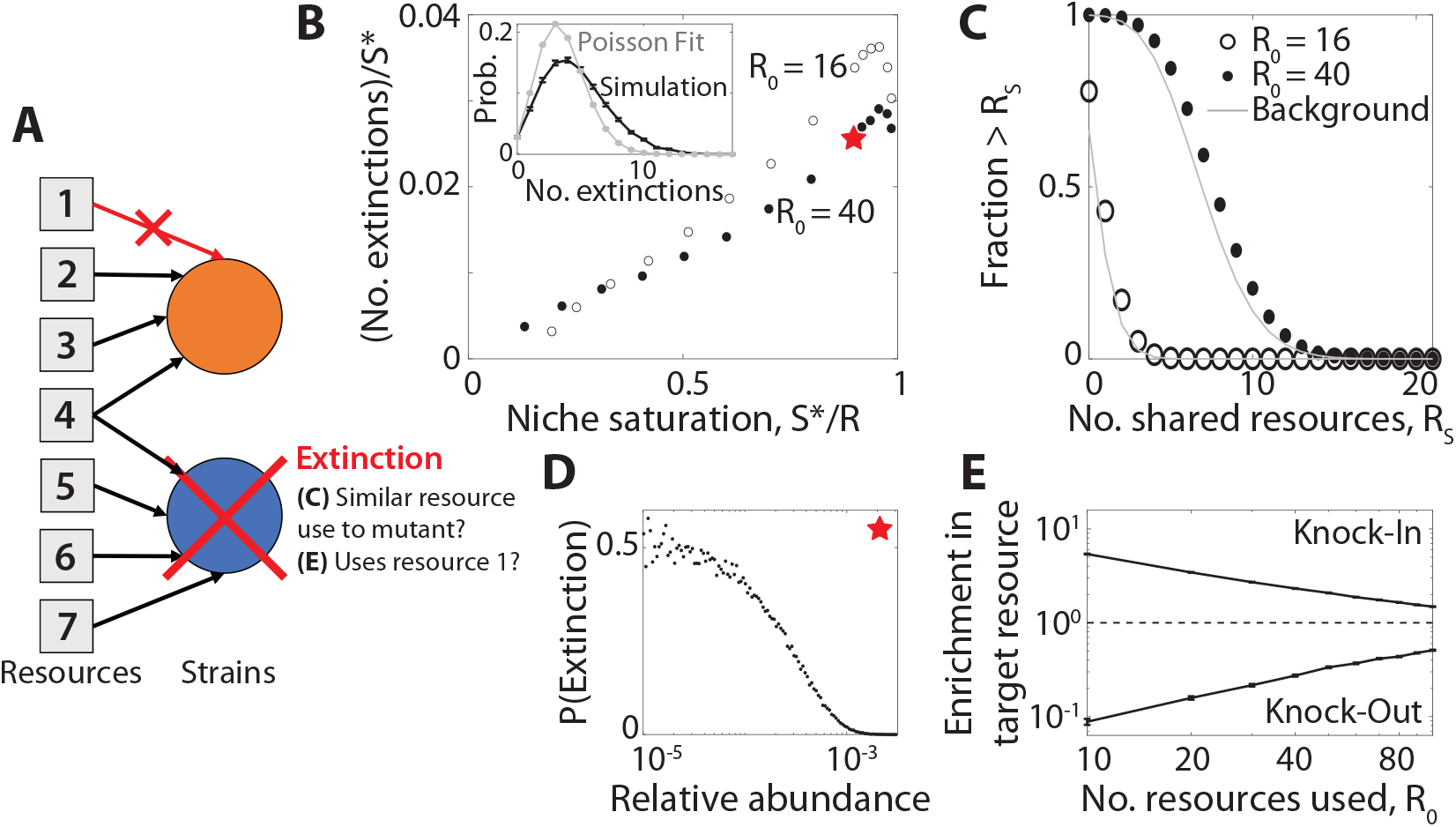
Successful mutations drive extinctions of metabolically distant species. (**A**) Schematic showing the extinction of an unrelated species (blue) after a beneficial knockout mutation (orange) invades. In this example, the displaced species and the mutant share one common resource, but not the one targeted by the knock-out mutation. (**B**) Average number of extinctions after the invasion of a successful mutant, as a function of niche saturation 𝒮^*^/ℛ ; parent strains are excluded from the extinction tally. Inset: full distribution of extinctions for the starred parameters, compared to a Poisson distribution with the same fraction of zero counts. Points denote the averages over 10^4^ simulation runs with the same base parameters as Fig. 3. (**C**) Distribution of the number of resources jointly utilized by the displaced species and the invading mutant (parent strains excluded). Points denote the results of simulations with 𝒮^*^/ℛ = 0.9. Gray curves show the analogous background distribution between the mutant and all other species in the community, regardless of whether they become extinct. (**D**) Probability of extinction as a function of initial relative abundance for the starred point in panel B. (**E**) The fold change in probability that the displaced species uses the same resource targeted by the mutant (i.e., the resource being knocked in or out), relative to the background distribution of resource use in the population. Points denote means and standard errors for knock-in and knock-out mutations as a function of ℛ_0_ for 𝒮^*^/ℛ = 0.8, with the the remaining parameters the same as panel B.

We tested this idea by examining the number of resources that were jointly consumed by the invading and displaced strains (a proxy for their overall metabolic similarity). Interestingly, we found that the number of resources shared with the mutant was not substantially higher for the displaced species, and was comparable to a randomly drawn species from the larger community (Fig. 4C). This suggests that the extinction events in Fig. 4 cannot be explained by traditional measures of niche overlap (33). Rather, successful mutants can displace species outside of their apparent niche, even when they stably coexist with a more metabolically similar parent.

Since extinctions were not well-predicted by their overall metabolic similarity to the mutant, we conjectured that these displaced species may possess other features that render them vulnerable to extinction. For example, low-abundance species may be less well-adapted to the current environment, and thus sensitive to perturbations like the invasion of a mutant. Consistent with this hypothesis, we found that the extinction probability for very low-abundance species is much higher than the community average, approaching ∼ 50% at the highest levels of niche saturation (Fig. 4D). Furthermore, although the displaced species are metabolically distant from the invading mutant, we find they are more likely to share the mutant’s strategy for the resource targeted by the mutation. Displaced species are more likely to use the resource gained by a successful “knock-in” mutation, and are correspondingly less likely to use the resource lost in a successful knock-out strain (Fig. 4E). These results illustrate that in high-dimensional ecosystems, the invasion of a new mutant can have a small impact on a diverse range of metabolic strategies. If a resident strain is maladapted enough to already be on the edge of extinction, the invasion of a mutant can be enough of a perturbation to displace it from the community. Thus, the abundance of an organism might often be a better predictor of its fate than its apparent metabolic niche.

### Robustness of ecosystem to subsequent mutations

Our analysis above focused on the first wave of mutations arising in a newly assembled community, where the initial states could be predicted using existing community assembly theory (28, 29, 35, 36). However, subsequent waves of evolution could eventually drive the ecosystem away from this well-characterized initial state (30). To assess the robustness of our results under the acquisition of further mutations, we simulated successive waves of mutations using a generalization of the approach in Fig. 3. We considered the simplest case where resident strains could generate knock-out mutations on any of the resources they currently utilized. We also assumed that the dynamics were mutation-limited (30, 31), so that the community relaxes to a well-defined steady state between each successive mutation. We continued this process until one of the surviving strains had accumulated 10 mutations in total, which typically corresponded to 100-200 successful mutations in the larger community.

We first asked how these subsequent waves of mutations altered the genetic structure of their community. While the total number of surviving strains decreased slightly over time (eventually stabilizing at an intermediate value), the number of strains related through *in situ* diversification events increased approximately linearly over the same time window (Fig. 5B). This indicates that the ecological diversification events in Fig. 3 continue to occur at a high rate even after additional mutations have accumulated, more quickly than individual branches of diversified lineages go extinct. Nonetheless, the abundance trajectories in Figure 5A indicate that these extinction events among close relatives are not uncommon. For example, Lineage 1 seeds eight ecological diversification events over the course of the simulation, but only three of these strains are alive at the end of the simulation. On average, we find that newly diversified lineages coexist for ∼ 40 mutational steps before one of them goes extinct (Fig. 5C). Nonetheless, longer coexistence is possible: one of the diversification events in Fig. 5A is maintained for *>*150 mutational steps, enough time for multiple additional mutations to accumulate within each lineage. Thus, coexisting mutant-parent pairs are often maintained through further evolutionary perturbations, even as the total number of species stabilizes over time.

**Figure 5:**
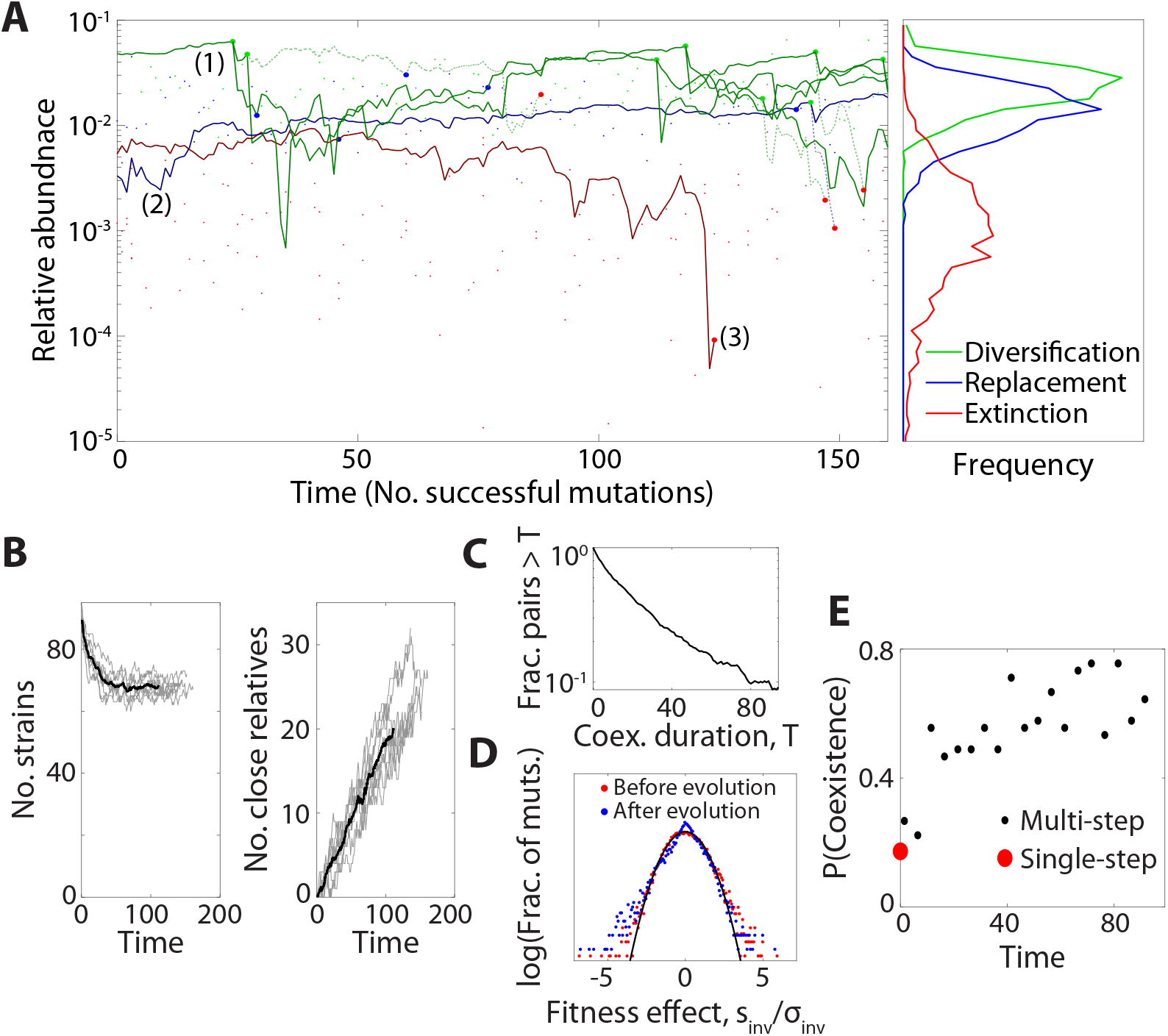
Mutation and diversification over longer evolutionary timescales. (**A**) Left: an example simulation showing the step-wise accumulation of ∼ 100 adaptive knock-out mutations in a community with ℛ= 100, ℛ_0_ = 20, 𝒮^*^/ℛ= 0.9, 𝒮^*^ /𝒮= 0.1. Solid lines denote the abundance trajectories of 3 example lineages. Large points indicate extinction events in these lineages (red), diversification events (green), and mutation events that displace their parent strain (blue) in the highlighted lineages, while smaller points indicate analogous events for other species in the community. Dashed lines illustrate offshoots of the highlighted lineages that eventually went extinct. Right: Relative abundances of strains when they experienced mutation, diversification and extinction events, respectively. Lines denote histograms aggregated over 9 simulation runs. (**B**) Left: Total number of surviving strains over time. Grey lines denote replicate simulations for the same parameters in panel A, while their average is shown in black. Right: Total number of strains related to another surviving strain through one or more *in situ* diversification events. (**C**) The probability of mutant-parent coexistence being maintained (i.e., both lineages surviving) as a function of time since initial divergence, over ten simulations. (**D**) The distribution of fitness effects at the start of the simulation (red) and after 90 accumulated mutations (blue), compared to the first-step predictions from Fig. 2 (black). (**E**) The probability that a mutation event leads to stable diversification as a function of time. Black points denote binned values aggregated over nine replicate simulations, while red point denotes the analogous result for 10^4^ first-step simulations.

Our first-step analysis showed that mutations in more abundant strains were more likely to coexist with their parent. This prediction is borne out by our multi-step simulations as well: while some highly abundant lineages seeded many *in situ* ecological diversification events (e.g. Lineage 1 in Fig. 5A), the typical surviving lineage experienced no more than one (e.g. Lineage 2), and many species went extinct without diversifying at all (e.g. Lineage 3). Consistent with our earlier predictions, lineages at low abundance were more prone to extinction when a new mutant invaded, while coexistence events tended to happen to high-abundance lineages (Figure 5A, right). The combination of these factors creates a “rich get richer” effect where high-abundance species diversify at the expense of driving low-abundance species to extinction (30).

Finally, we asked how the accumulation of mutations shifts the landscape of beneficial mutations from the initial “assembled” state in Figs. 2 and 3. We found that the distribution of fitness effects does not dramatically differ between the start and the end of the simulation, although its shape changes slightly from a Gaussian distribution toward a two-tailed exponential (Fig. 5D). In contrast, the impacts of successful mutations exhibit larger changes: the proportion of mutation events which result in mutant-parent coexistence increases substantially with evolutionary time, rising from about 20% to 80% over the course of the simulation (Fig. 5E). This shift cannot be explained by changes in the overall number of species or the availability of beneficial mutations, but instead deviates entirely from the replica-theoretic predictions that describe our initial state. Thus, while many our qualitative conclusions continue to hold on longer evolutionary timescales, the accumulation of mutations can lead to quantitative differences from our first-step analysis above. In both cases, the evolved communities exhibit distinct genetic signatures compared to their purely assembled counterparts, which could motivate future tests of *in situ* evolution.

## Discussion

Large ecological communities apply complex evolutionary pressures to their resident species, leading to controversy about how these organisms’ evolutionary trajectories are affected by the background community (17). Here, we address this challenge by developing a theoretical framework for predicting the first steps of evolution in large, randomly assembled communities that compete for substitutable resources. These “mean-field” results hold when surviving strains consume many resources, and each resource is consumed by many surviving strains. Under these conditions, our results provide a mechanistic approach for understanding how the fitness benefits and fates of new mutations should scale with the diversity and metabolic overlap of the surrounding community.

Our results show that the supply of beneficial mutations does not necessarily run out in larger communities – as expected in the simplest models of niche filling – but rather that the benefits of these mutations will systematically decline with the degree of niche saturation (𝒮^*^/ℛ). We also find that the fitness effects of mutations can be broadly correlated with the external environment, even in large communities where internal resource concentrations are broadly shielded from external environmental shifts (32, 36). These distributions of fitness effects can be measured in modern experiments, e.g. by performing barcoded fitness assays in communities of different size (12, 50).

Our finding that successful mutants often coexist with their parents is reminiscent of empirical observations from the gut microbiome, where recently diverged strains differing by only a handful of mutations appear to stably coexist within their host (5, 6). While spatial structure could also contribute to this coexistence (27, 53), our model demonstrates that *in situ* diversification can continue to occur even in a well-mixed environment when most niches are already filled. Similar behavior has also been observed in generalized Lotka-Volterra models with spatiotemporal chaos (54). This suggests that ongoing diversification may be a generic feature of large microbial communities, providing an alternative mechanism for the “diversity begets diversity” effect (17, 55, 56) that does not rely on explicit cross-feeding interactions.

Beyond diversification, Figure 5 demonstrates that successful mutants can also drive distantly related species to extinction. This behavior is consistent with recent empirical observations in the human gut microbiome (8) and strain-swapping experiments in synthetic gut communities (57). It could also contribute to the extinction events observed in community passaging experiments (58). The fact that the displaced species are metabolically diverged from the invading mutants creates obstacles for inferring these interactions from metabolomics measurements (59) or metabolic reconstructions (60). Our findings suggest that future efforts should instead focus on how the invading mutant impacts low-abundance species more generally, and whether they utilize the specific resource(s) targeted by the mutation. These results align with recent work emphasizing the importance of collective interactions in shaping microbial community dynamics (61).

Here, we focused on the simplest possible resource competition model, neglecting important factors like spatial structure (62), metabolic regulation (46), and more general forms of cross-feeding (49, 63), which are all thought to play key roles in natural microbial communities. We also considered the simplest regime of evolutionary dynamics that neglect competition between simultaneously occurring mutations. These clonal interference effects can enhance diversity by allowing strains to temporarily evade competitive exclusion (31). On the other hand, competition between lineages also selects for more strongly beneficial mutations (64), which we predict are less likely to coexist with their parent. Our results provide a baseline for incorporating these effects, which will be crucial for understanding how large microbial communities will evolve.

## Author contributions

Conceptualization: J.M. and B.H.G.; theory and methods development: J.M. and B.H.G..; analysis: J.M. and B.H.G.; writing: J.M. and B.H.G.

## Code Availability

Source code for community assembly simulations, numerical calculations and figure generation are available at Github (https://github.com/jdmcenany/First_Step_Muts).

## Supplementary Information

**Figure S1:**
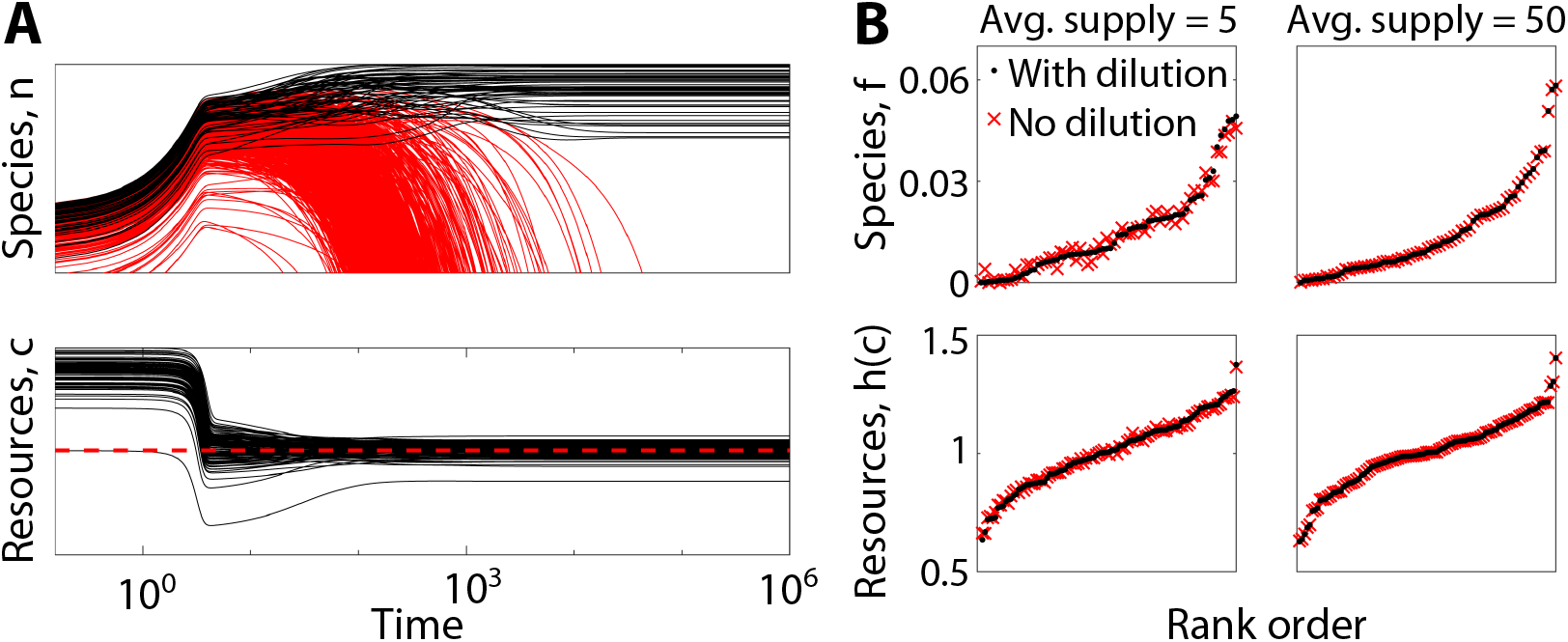
**(A)** Absolute abundance of species (top) and resource concentrations (bottom) over time on a log-log scale, for an example community simulated by explicitly integrating Eq. (S1a). Red curves are species which go extinct at equilibrium; the red dashed line in the lower plot shows the expected average value of *c* at equilibrium. **(B)** Equilibrium species relative abundances and resource availabilities *h*(*c*), obtained from explicit simulation (black dots) and finding the equilibrium through optimization, ignoring the dilution term (red crosses). Predictions are shown for two values of average resource supply 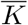. Simulations were performed with ℛ = 100, ℛ_0_ = 20, 𝒮/ℛ = 10, Std(*X*_*µ*_) = 0.5/ℛ_0_, and 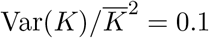. The uptake function *u*(*c*) was taken as a Monod function, *u*_*i*_(*c*_*i*_) = *u*_max_*c*_*i*_/(*c*_1/2_ + *c*_*i*_), with *u*_max_ = 4 and *c*_1/2_ = 1.

**Figure S2:**
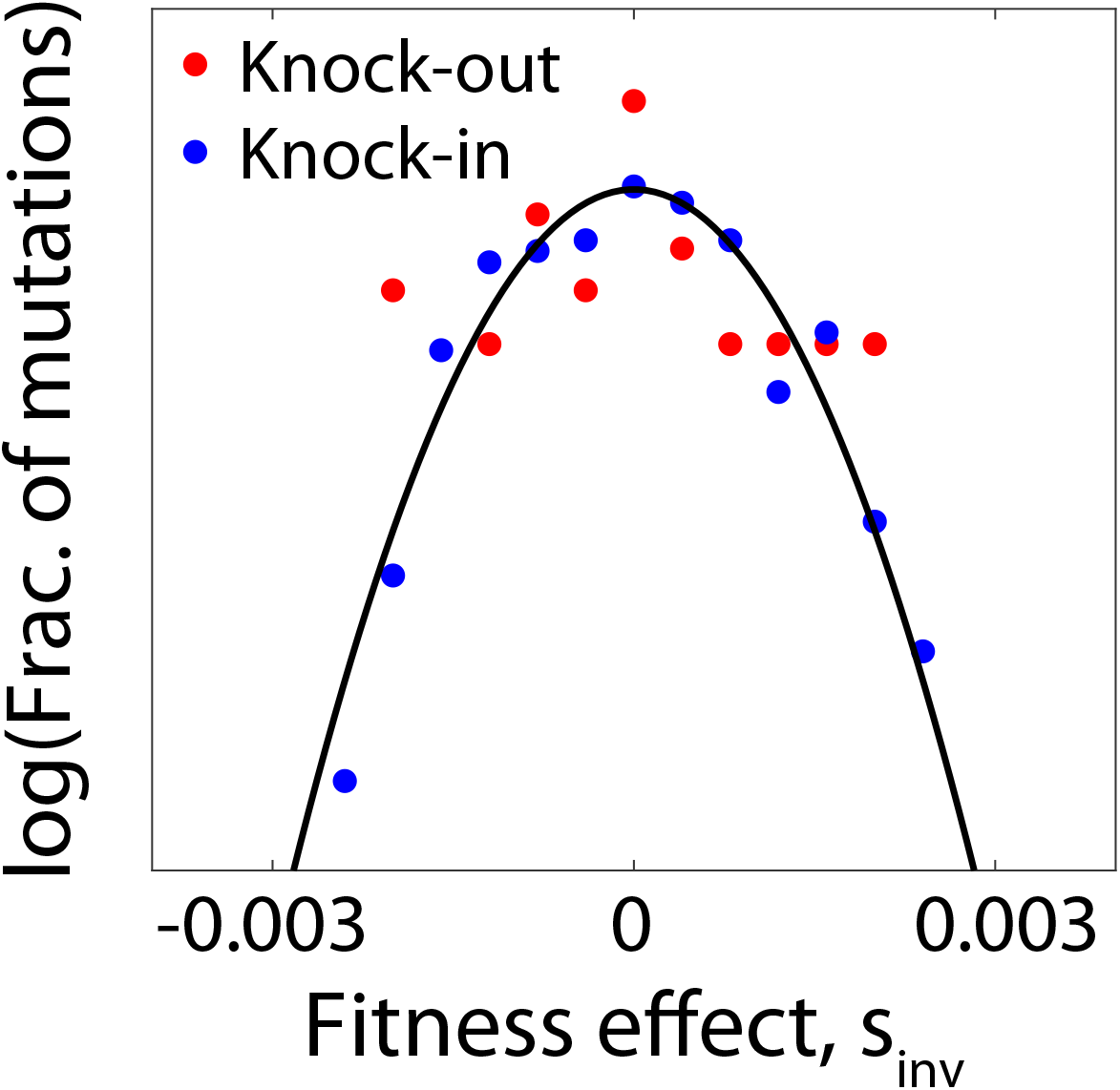
Distribution of invasion fitnesses for knock-out and knock-in strategy mutations of a single organism within a single sampled community. Black curve shows Gaussian theory prediction, while dots are histogram values over all possible strategy mutations. Simulation was run for 𝒮^*^/ℛ = 0.8, *ℛ*= 200, ℛ_0_ = 40, 𝒮^*^/𝒮 = 0.1, and uniform resource supply.

**Figure S3:**
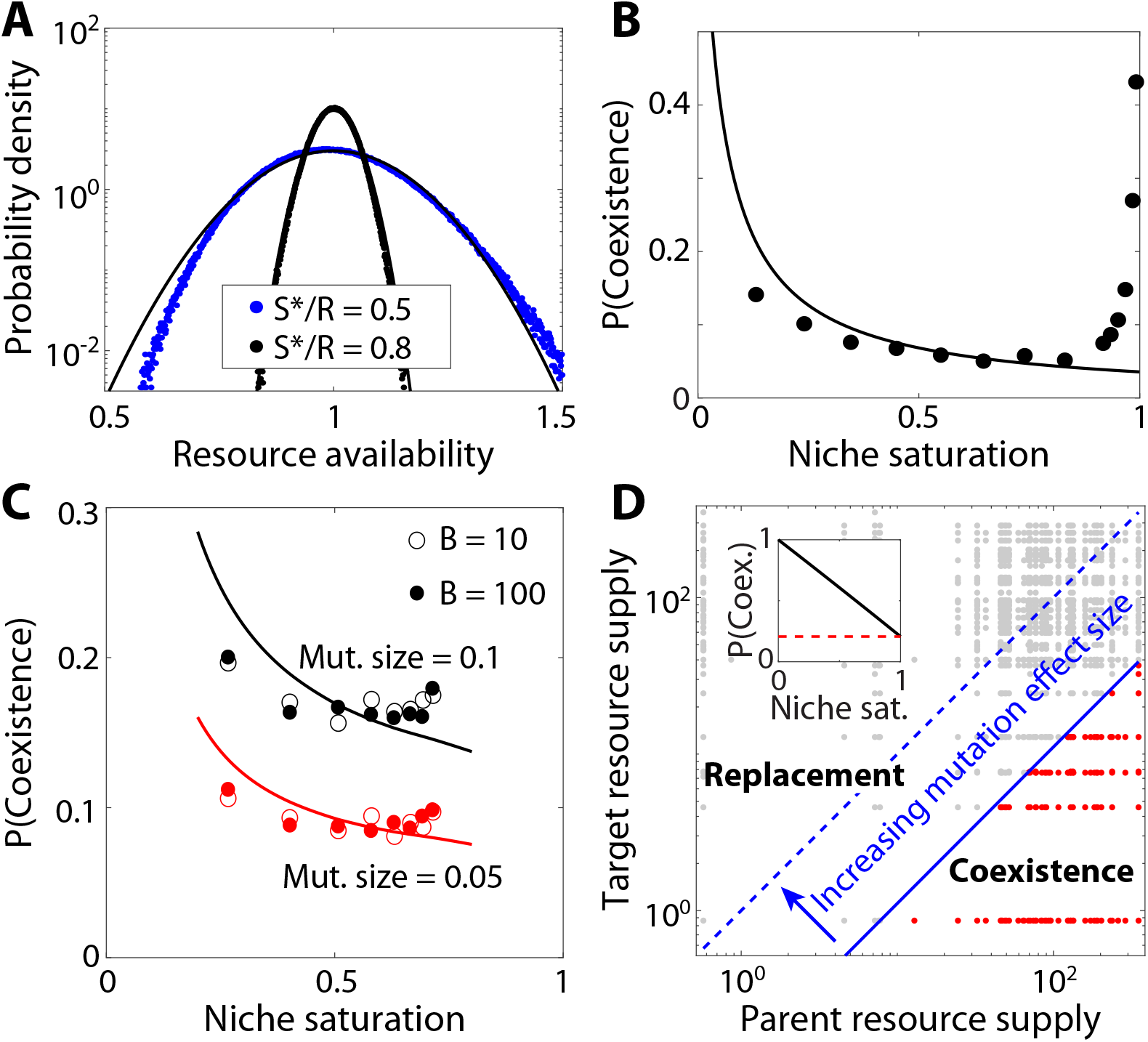
Our main conclusions about mutation in assembled communities can be generalized to other consumer-resource sampling structures. **(A)** Distribution of resource availabilities *h*_*i*_ for an ecosystem assembled with strategy vectors drawn from the Dirichlet distribution as described in SI Section 4.4. Black curves show theoretical prediction from the binary resource use case studied in this paper, while points show simulated results. Simulations were run with ℛ= 200, ℛ_0_ = 40 and 𝒮^*^/𝒮 = 0.1 for uniform resource supply. **(B)** Probability of mutant-parent coexistence for knock-out strategy mutations in this Dirichlet assembled community, with the same parameters as in (A). Black curve shows the theoretical prediction for binary resource use. **(C)** Mutant-parent coexistence probability for a community where all species consume all resources at similar levels (SI Section 4.4), for different mutation effect sizes Γ. Curves show theoretical prediction from the binary model, with an effective value of ℛ_0_ corresponding to the shape parameter *B* of the Dirichlet distribution. Other parameters were Std(*X*_*µ*_) = 1/*Bℛ* and ℛ= 200. **(D)** Conditions for mutant-parent coexistence in a specialist community (SI Section 4.5). Each point corresponds to a beneficial mutation in a 50-member community, with red points showing mutant-parent coexistence. Axes indicate the supply of the resource the parent consumes, and the supply of the “target” resource the mutant gains access to. The blue line is the predicted boundary between coexistence and replacement for mutations with effect size γ_spec_ = 0.1. Inset: Mutant-parent coexistence probability as a function of niche saturation; the red dashed line corresponds to a saturated community, as shown in the rest of (D).

**Figure S4:**
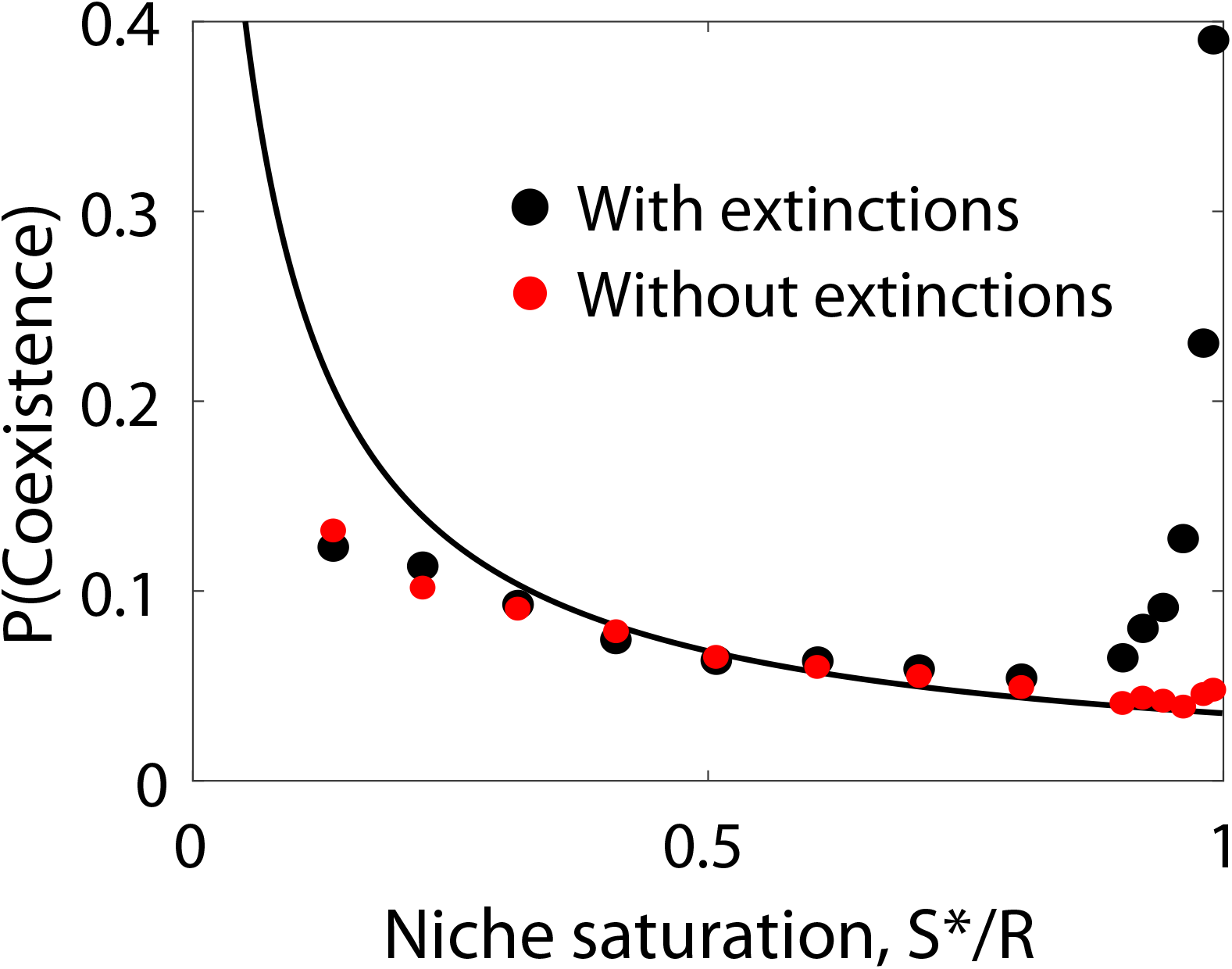
Probability of coexistence as a function of niche saturation, with simulations using the “simultaneous assembly” approximation where mutant invasion is modeled using a single assembly process (SI Section 2.1). Red points show the simultaneous assembly approximation, which allows reappearance of extinct species. Black points show simulations where extinct species were disallowed from returning (as in the rest of our study); this change results in coexistence probability increasing at high saturation. Simulations were performed with ℛ = 200, ℛ_0_ = 40, and 𝒮^*^/𝒮 = 0.1 using knock-out mutations.

**Figure S5:**
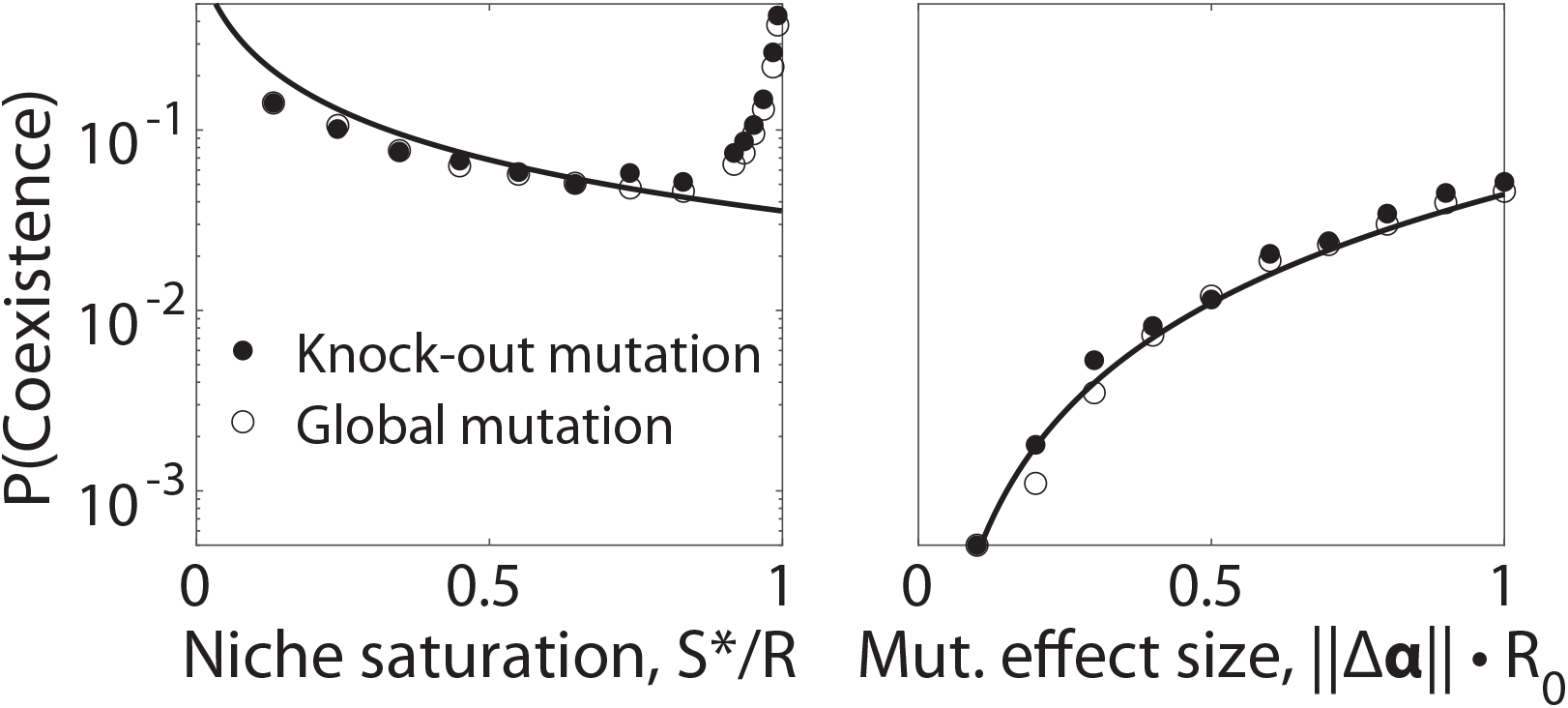
Coexistence probability for different strategy mutations. Left: Mutant-parent coexistence probability as a function of niche saturation 𝒮^*^/ℛ for full knock-out mutations and “global” strategy mutations with 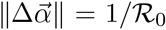 (which have similarly-sized changes to all nonzero entries of 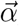, as described in SI Section 4.4). Right: Mutant-parent coexistence probability as a function of mutation effect size 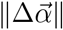 for 𝒮^*^/ℛ = 0.8, for partial knock-out and global strategy mutations. Black curves show theoretical prediction. All simulations run with ℛ= 100, ℛ_0_ = 40, and 𝒮^*^/𝒮 = 0.1 using Dirichlet-distributed resource usage vectors.

**Figure S6:**
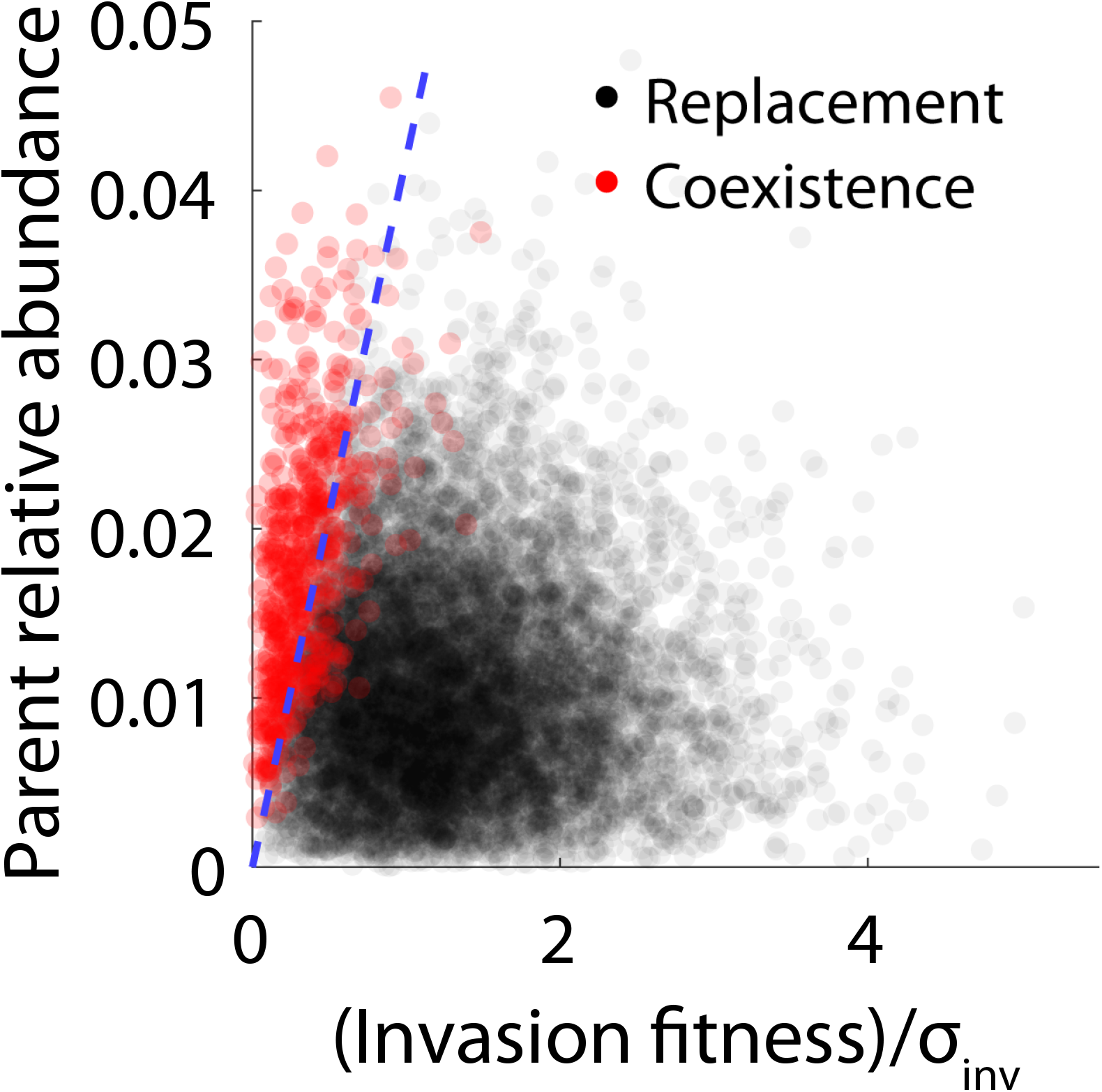
Scatterplot showing the invasion fitness of knock-out strategy mutations against the relative abundance of the parent before mutant invasion, with colors indicating whether the mutants coexist with or replace their parent at ecological equilibrium. The dashed blue line shows the theoretical prediction that should divide replacement events from coexistence events. Simulations run for ℛ = 200, ℛ_0_ = 40, 𝒮^*^/𝒮 = 0.1, and 𝒮^*^/ℛ = 0.8.

**Figure S7:**
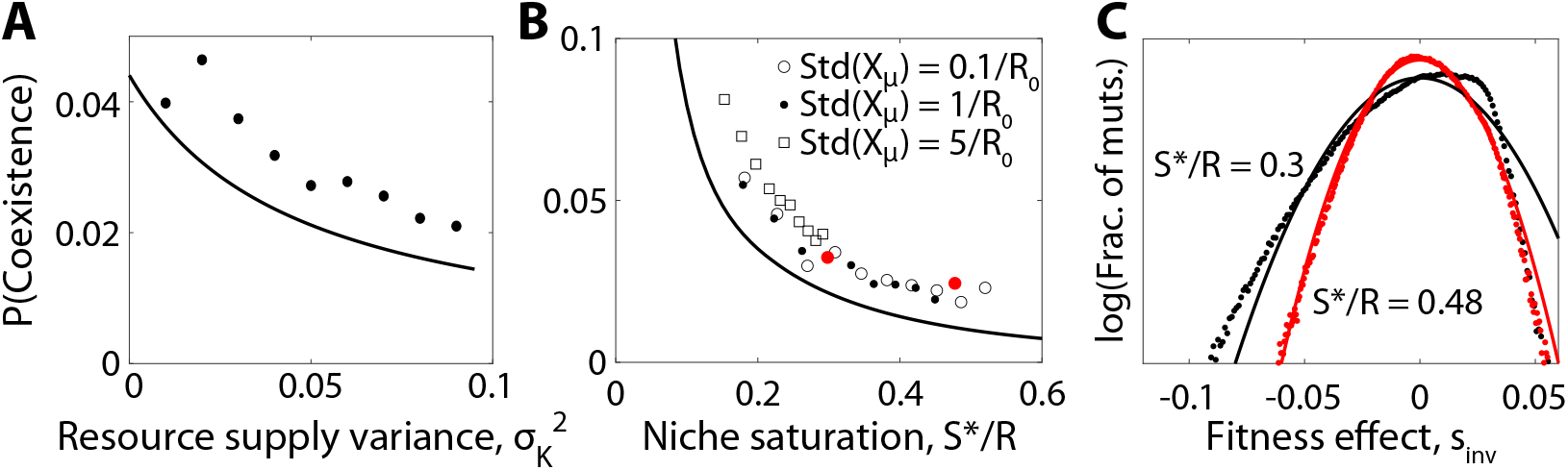
The distribution of fitness effects and mutant-parent coexistence probability in the presence of heterogeneous resource supply rates. **(A)** Mutant-parent coexistence probability for knockout mutants as a function of scaled variance in resource supply 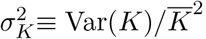, with ℛ = 200, ℛ_0_ = 40, 𝒮^*^/𝒮 = 0.1, and 𝒮^*^/ℛ= 0.8. **(B)** Coexistence probability of parents and knock-out mutants for communities assembled with exponentially distributed resource supply (corresponding to 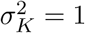), and different levels of uptake budget variation Std(*X*_*µ*_) among sampled organisms. Simulations were run with ℛ= 150 and ℛ_0_ = 30. **(C)** Distribution of fitness effects for knock-out mutants for communities corresponding to the red points in (B). In all plots, points show simulation results while curves show theory predictions.

**Figure S8:**
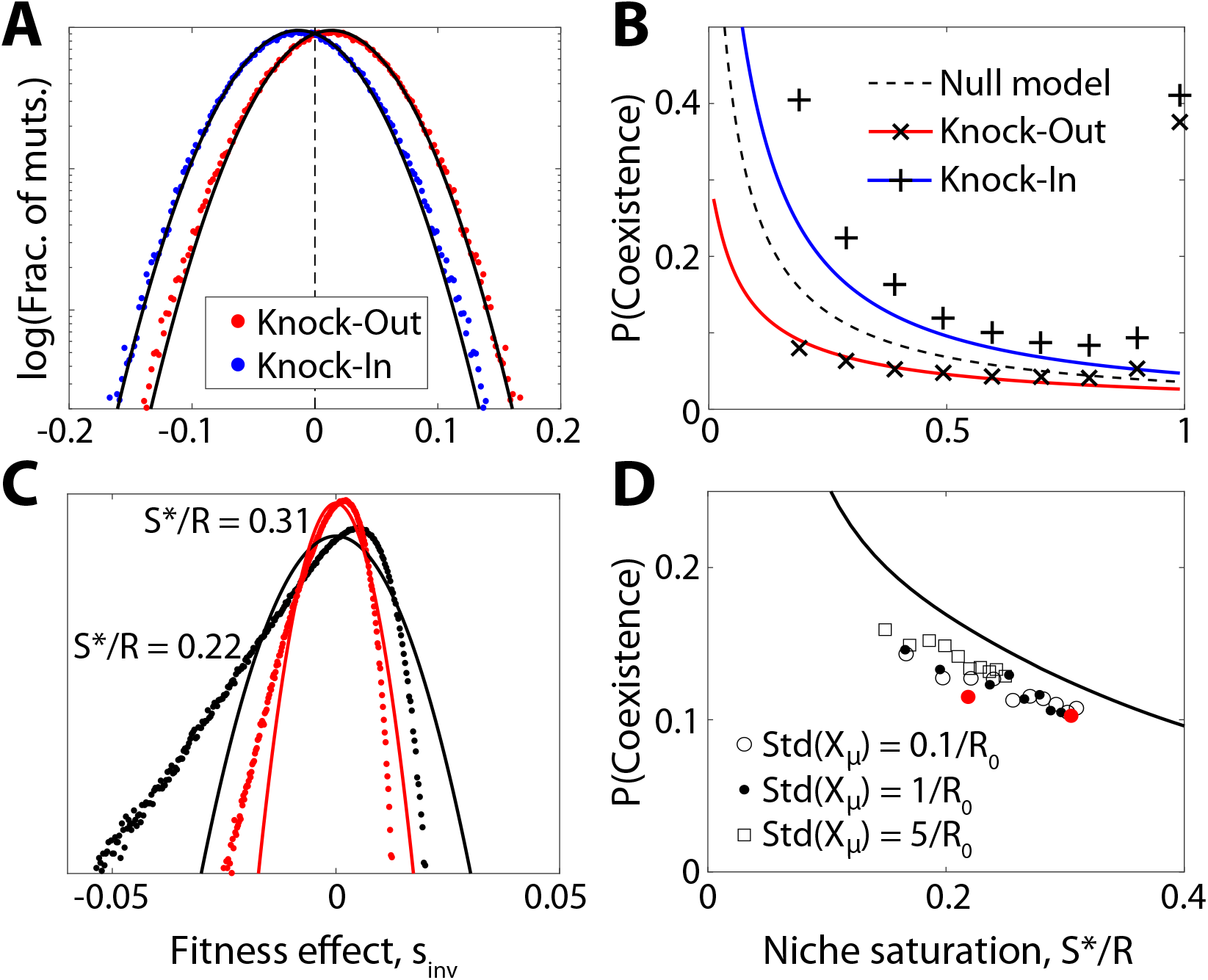
**(A)** Distribution of invasion fitnesses for the alternative model in Ref. (36), where the cost of using a resource contributes to the death rate of an organism, rather than occupying a portion of its overall resource uptake budget. Black curves show theoretical prediction for knock-in and knock-out mutations, while points show simulation results. Simulations run with ℛ= 200, ℛ_0_ = 40, 𝒮^*^ /𝒮 = 0.1, and 𝒮^*^/ℛ = 0.8. **(B)** Mutant-parent coexistence probability for the Ref. (36) CRM with the same parameters as (A), as a function of niche saturation, 𝒮^*^/ℛ. The average fitness benefit (cost) for knock-out (knock-in) mutations effectively gives strategy mutations a nonzero change in pure fitness, which lowers (raises) the theoretically predicted coexistence probability relative to the neutral case. Red and blue curves show these adjusted predictions, while crosses and pluses show simulated results. **(C)** Distribution of fitness effects for knock-out mutations in a community where sampled species lack metabolic trade-offs, as described in SI Section 4.3. Simulations run with ℛ= 150 and ℛ_0_ = 30. **(D)** Mutant-parent coexistence probability for the same ecosystem as a function of niche saturation, for different levels of variation in uptake budget Std(*X*_*µ*_). Red points correspond to the parameter values in (C).

**Figure S9:**
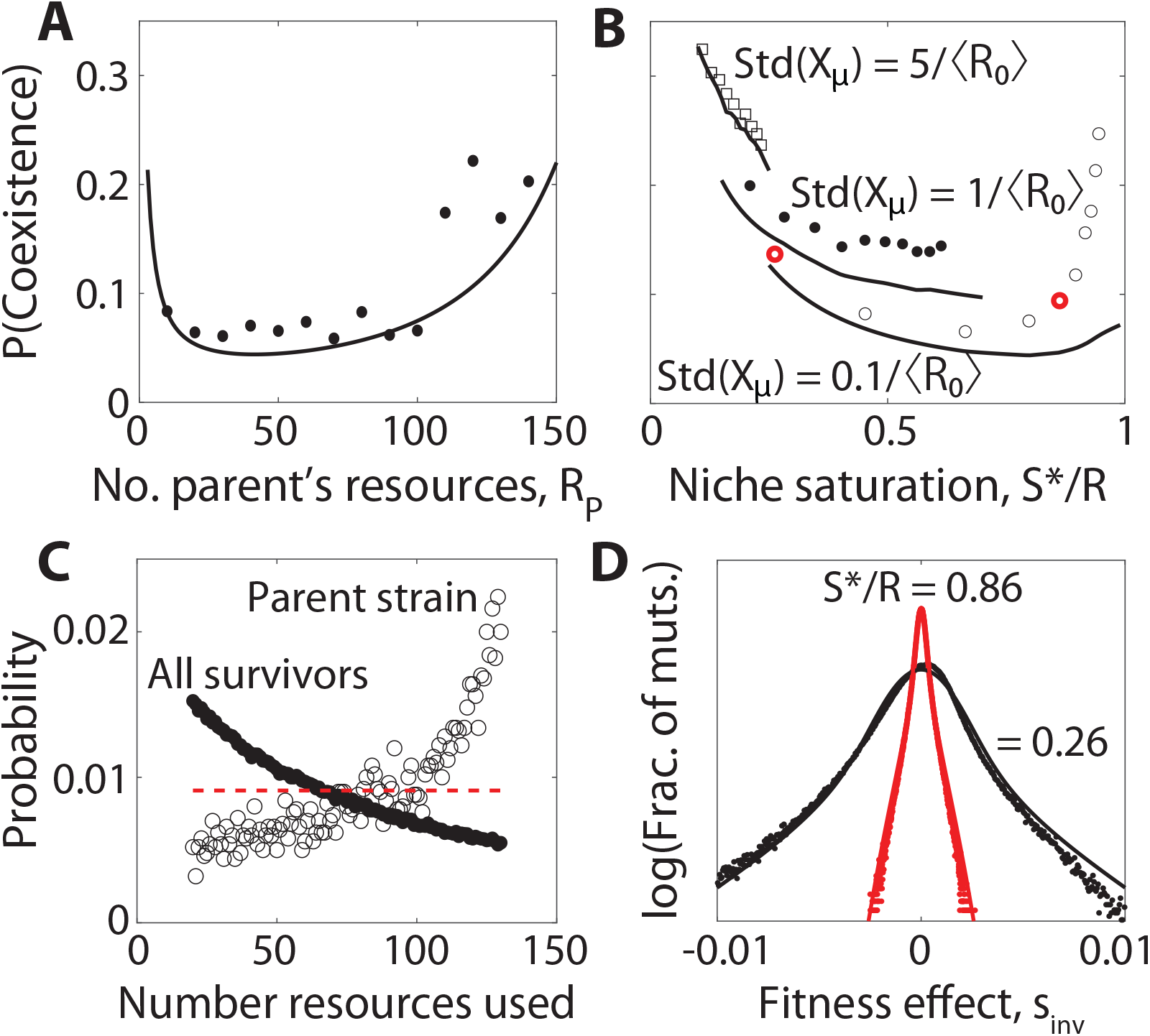
Extensions to communities with wider variation in the total number of resources consumed per species. **(A)** Probability that the mutant coexists with its parent strain as a function of the number of resources ℛ_*P*_ used by the parent, in a background community with ℛ_0_ = 40. Points denote simulation results for ℛ= 200, 𝒮^*^/𝒮 = 0.1, and 𝒮^*^/ℛ = 0.8. **(B)** Mutant-parent coexistence probability when the number of resources used by each strain is uniformly distributed between 30 and 120 (SI Section 4.2). Points denote simulation results for knock-out mutations with ℛ= 150, and different levels of variation in uptake budget Std(*X*_*µ*_). **(C)** Distribution of the actual number of resources used by surviving species in the community corresponding to the rightmost red circle in (B), including all survivors (closed circles) and the subset of species which produced a successful knock-out mutation (open circles). The red dashed line shows the corresponding uniform distribution for sampled species. **(D)** Distribution of knock-out fitness effects for the community in (B), with parameter values corresponding to the red circles. All points show simulation results, while curves show theory predictions, using the distribution in (C) as an input.

**Figure S10:**
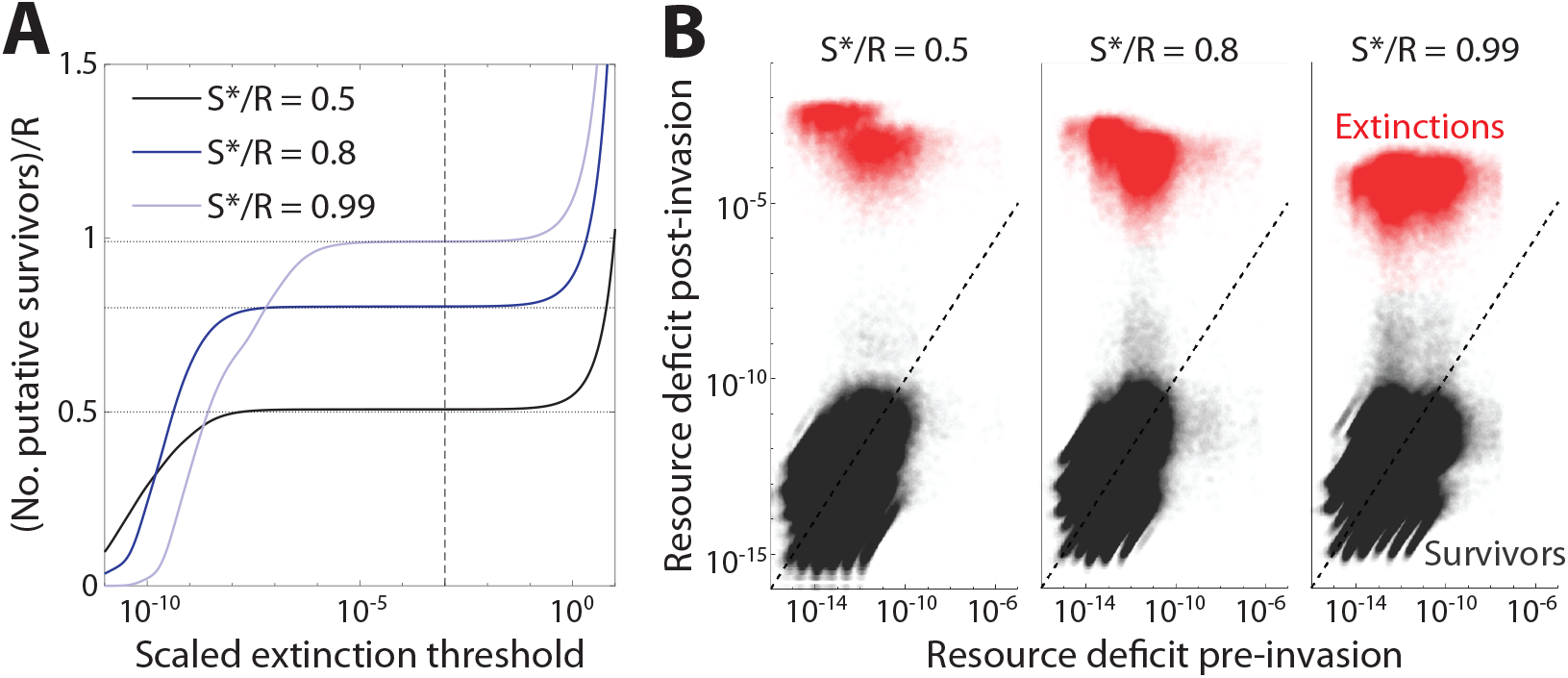
Checks to ensure that we have consistently identified extinctions numerically. **(A)** Plot of the number of putatively identified survivors for an assembled community, as a function of the numerical extinction threshold. If the resource deficit for an organism is above this threshold times Std(*h*_*i*_), the organism is labeled extinct. The plateau over multiple orders of magnitude indicates a numerically stable definition of survival vs. extinction; the dashed line shows the threshold used throughout our analysis. **(B)** Scatterplot of numerical resource deficit of organisms before and after invasion of a knock-out strategy mutant, with color indicating whether the resource deficit after invasion passes the extinction threshold, (i.e., whether the species has gone extinct). The fact that most red and gray points are clearly separated shows that most species identified as extinct are insensitive to our precise choice of threshold. Simulations were run with ℛ = 200, ℛ_0_ = 40, 𝒮^*^/𝒮 = 0.1.

### 1 Model Outline

In this section, we outline the setup that allows us to derive our main theoretical results. First, we describe the consumer resource model we study, and how its dynamics can be coarse-grained to a limit where species consume most of the available resources. The task of finding an ecological equilibrium in this coarse-grained model can be described as an optimization problem. Then, we define a convenient fitness gauge which brings our model in line with previous work.

#### 1.1 General Ecological Dynamics

As summarized in the Model section, we are interested in the following system: consider an ecosystem formed by combining 𝒮 species, identically and independently drawn from some underlying distribution, competing for ℛ resources. We allow this ecosystem to come to ecological equilibrium, at which point 𝒮^*^ ≤ 𝒮 species will survive at nonzero abundance. Then, suppose that one randomly selected surviving species undergoes a mutation to produce a new strain, which can differ from its parent in its ability to consume resources. If the mutant has positive invasion fitness, it will rise to nonzero abundance in the population, and the abundances of other species may change as well. We are interested in analyzing the properties of this new ecosystem with a single mutant, particularly the probability that the mutant and its parent strain are able to coexist at ecological equilibrium.

First, we define our community assembly process and consumer resource model, which generally follows previous work (28–32, 36, 45). We consider a well-mixed community formed from the combination of 𝒮 strains, indexed by *µ* ∈ {1, …, *𝒮*}, and ℛ resources, indexed by *i* ∈ {1, …, ℛ}. Each organism is associated with a resource uptake vector,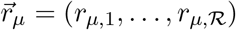, which could represent the expression of enzymes used to metabolize each resource. The consumer-resource dynamics for the strain abundances (*n*_*µ*_) and the local resource concentrations (*c*_*i*_) are given by Eq. (1) in the Main Text:

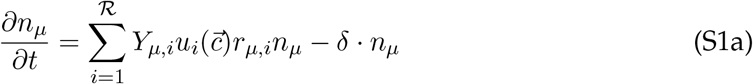

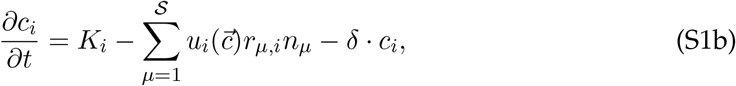

where *δ* is the dilution rate, *K*_*i*_ is the input flux of each resource, 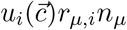 is the total uptake rate of resource *i* by strain *µ*, and *Y*_*µi*_ is the yield of resource *i* when converted into biomass by strain *µ*. For simplicity, we will assume that this yield matrix can be factored into a rank-one form *Y*_*µ,i*_ = *a*_*i*_/*b*_*µ*_. While this assumption can approximately account for effects like different stoichiometric yields and cell sizes, it neglects idiosyncratic effects such as resource sequestration (47), where some organisms remove resources from the environment without corresponding growth in abundance. This simplification allows us to choose “biomass-equivalent” units such that *Y*_*µ,i*_ = 1, yielding a conservation of mass equation corresponding to a symmetry between resource consumption and biomass growth:

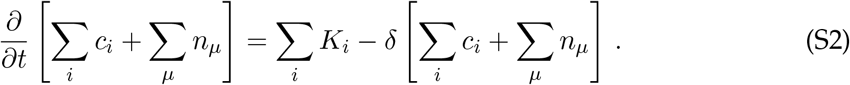

The total mass rapidly approaches the 𝒮 + ℛ − 1 dimensional manifold,

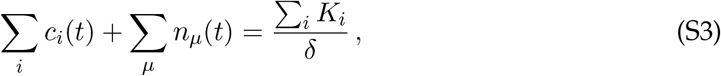

on a timescale proportional to the inverse dilution rate *δ*^−1^ (which is equivalent to the average generation time at steady state). Since we are interested in the dynamics on timescales that are much longer than a single generation, we assume that Eq. (S3) always applies.

#### 1.2 Coarse-Grained Ecological Dynamics

The dynamics in Eq. (S1) can be further simplified if we assume that the resource uptake rates are sufficiently fast, and the total biomass of organisms sufficiently large, that most supplied resources are consumed by organisms rather than diluted out. When this is the case, we can coarse-grain over time such that the resource concentrations are instantaneously set by the abundance of each strain, rather than being independent dynamical variables (“integrating out the resources”). While this procedure follows previous work (30, 32), we will restate this approximation and its self-consistency conditions for completeness.

As shown by Equation (S1), it is reasonable to approximate that most resources are converted to biomass rather than diluted out when the local concentration of each resource is much smaller than 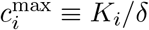, which is the equilibrium value that would be obtained in the absence of any microbes. In other words, the community must be able to deplete each resource far below the level that would be expected from the environmental supply rates alone. When this condition is satisfied, the total biomass in Eq. (S3) will be dominated by the microbial contribution,

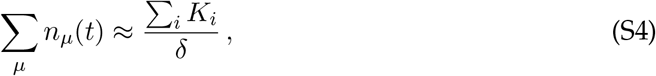

and the uptake rates of each resource will approach a local steady state set by the current strain abundances:

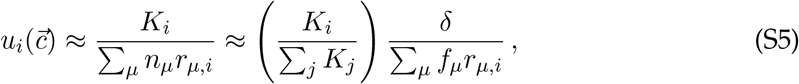

where we have defined the relative abundance variables,

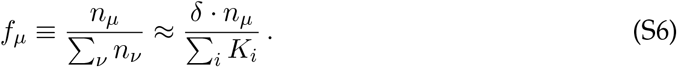

Substituting these expressions into Eq. (S1a) yields the coarse-grained model in Eq. (2) in the Main Text:

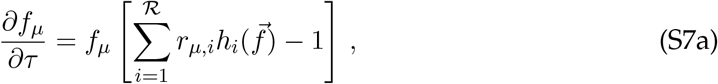

where time is measured in generations (τ ≡ δ · t) and

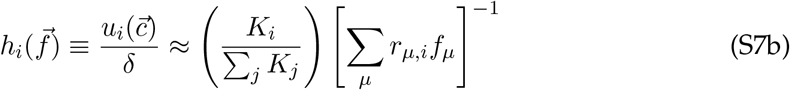

can be interpreted as the local availability of resource *i*.

##### Self-consistency of coarse-grained dynamics

These coarse-grained dynamics will be self-consistent when the relative abundances that are attained in Eq. (S7) satisfy

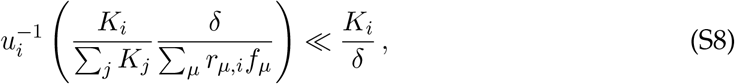

where 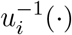 is the inverse of the uptake function 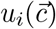. Since the relative abundances only depend on the relative supply rates, *K*_*i*_ /*∑*_*j*_ *K*_*j*_, this condition can always be satisfied if *δ* is sufficiently small, or if the total biomass is sufficiently large. For example, if the uptake rates can be described by a Monod function,

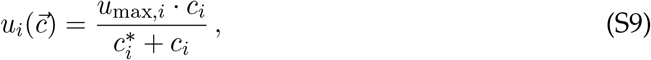

then the condition in Eq. (S8) reduces to

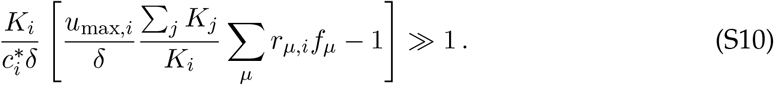

This will be valid for small enough values of *δ*, or large enough values of ∑_*j*_ *K*_*j*_/*K*_*i*_.

##### Rescaled variables

In our analysis below, it will be convenient to reparameterize the model in Eq. (S7) by decomposing the resource uptake vector 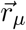 into an overall “uptake budget” *X*_*µ*_ ≡log ∑_*i*_ *r*_*µ,i*_, and a normalized “strategy vector”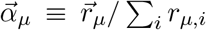, which contains ℛ non-negative entries that sum to one. The original uptake rates can then be expressed as

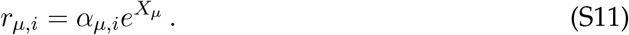

The uptake budget dictates the overall capacity of the organism to consume resources, setting its maximum growth rate under ideal growth conditions; the strategy vector describes how the organism allocates this uptake budget between the different substitutable resources. This alternative parametrization is equivalent to the original basis 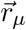, but will lead to simpler expressions in many of our analytical calculations below.

To streamline notation, it will also be useful to define the relative resource supply rates,

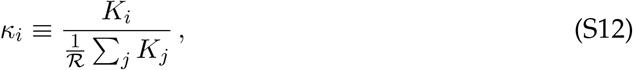

which describe how the supply of a given resource deviates from the average value, 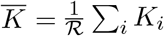. The relative supply rates are normalized such that 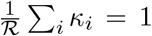. We will also refer to the variance of 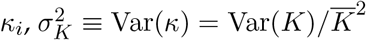.

#### 1.3 Incorporating Simple Forms of Cross-feeding

While the derivation of Eq. (S7) assumed that the resources were externally supplied, similar dynamics can continue to hold for simplified forms of cross-feeding, provided that *κ*_*i*_ is replaced by an effective supply rate that accounts for internal metabolic conversion. To see this, we consider the “universal” cross-feeding model from Ref. (49), where the metabolism of each resource is associated with a corresponding “leakage rate” *ℓ*_*i*_, which describes the proportion of that resource that is excreted back into the environment in the form of other resources, rather than being converted into biomass. This conversion process is described by a stoichiometric matrix **D**, whose entries *D*_*i,j*_ represent the fraction of resource *j* that is excreted in the form of resource *i*. This model assumes that every organism capable of metabolizing a resource produces the same byproducts in the same proportions; species can still differ in their overall excretion profiles, but these are completely determined by the corresponding uptake rates 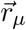. These cross-feeding dynamics can be described by a generalization of Eq. (S1),

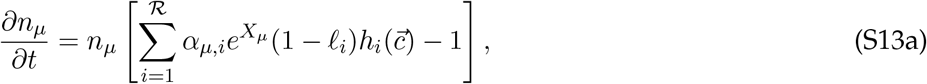

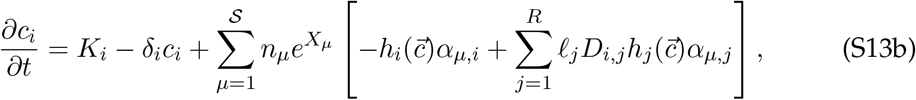

where the first term in the brackets of Eq. (S13b) represents the consumption of that resource, while the second term represents the production via metabolism of other resources. If we can continue to make the approximation that the dilution of resources is negligible compared to their consumption by other organisms, then the steady-state resource availabilities 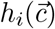 will obey a generalization of Eq. (S5),

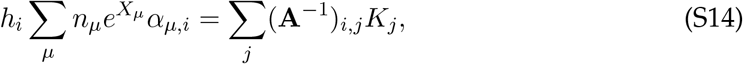

where **A** is an ℛ-by-ℛ matrix whose entries are given by *A*_*i,j*_ = *δ*_*i,j*_ − *ℓ*_*j*_*D*_*i,j*_ (note that *δ*_*i,j*_ is the Kronecker delta). This shows that we can recover the same dynamics in Eq. (S7) by defining the effective supply rates,

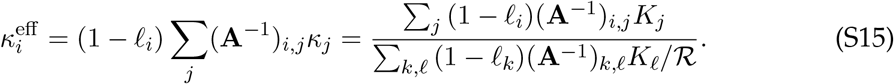

which depends on the external supply rates and the stoichiometry of metabolic conversions, but is otherwise independent of the species that are present in the community.

#### 1.4 Defining the Local Species Pool

All that remains is to choose the resource consumption parameters of the 𝒮 initial colonizers. As described in the text, we assume that the phenotypes of each species are drawn from a common statistical distribution. For our theoretical analysis, we follow the procedure of Ref. (36), where the uptake budgets *X*_*µ*_ are drawn from a normal distribution with standard deviation *ϵ*/ ℛ_0_, and the resource consumption strategies 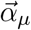 are normalized binary vectors where each entry has an independent probability ℛ_0_/ℛ of being nonzero. The independence of *X*_*µ*_ and 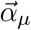 results in a soft metabolic tradeoff, where strains that consume more resources are usually less effective at using any individual one. To test the importance of this and other assumptions, we also perform simulations for several alternative distributions (such as Dirichlet-distributed resource consumption strategies) to verify that our qualitative results are robust (SI Section 4; Figures S3, S8, and S9).

#### 1.5 Ecological Equilibrium as Optimization

For any fixed set of species, the ecological dynamics in Eq. (S7) can be shown to possess a convex Lyapunov function,

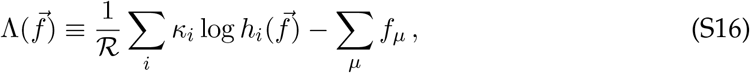

and a single stable equilibrium (see, e.g. Refs. 30, 32, 45). At this equilibrium, none of the surviving strains can increase in abundance, while none of the extinct strains could invade the ecosystem if they were re-introduced at low abundance. Thus, at ecological equilibrium, the bracketed term in Eq. (S7) (the “resource surplus”) must be non-positive for all *µ*, and is equal to zero if and only if species *µ* survives at nonzero abundance.

Conveniently, the task of finding the unique stable equilibrium in this model can be recast as a constrained optimization problem (30, 36, 37). Specifically, we treat the resource availabilities *h*_*i*_ (rather than the relative abundances *f*_*µ*_) as free variables to optimize over, and consider the task

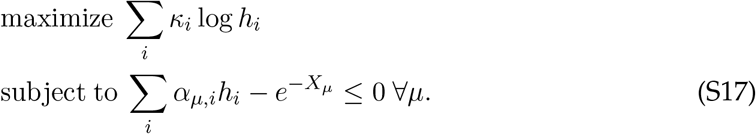

The constraint is the same as that discussed above, requiring that the resource surplus be non-positive for each organism. This model possesses an important symmetry which will be useful when making approximations later: it is invariant under a rescaling of the total uptake budget, which we can view as a change in “pure fitness” decoupled from the environment. In particular, if we change variables into translated fitness 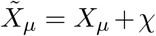, and rescaled 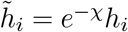, then the optimization problem becomes

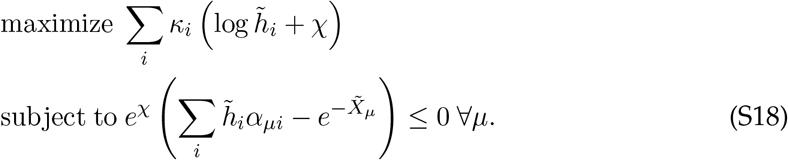

Since 𝒳 is just an arbitrary constant, it is clear that these optimization problems are the same, up to the rescaling of the *h*_*i*_. Suppose that 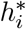 solves the first optimization problem. Then, 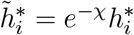 will solve the gauge-shifted optimization problem

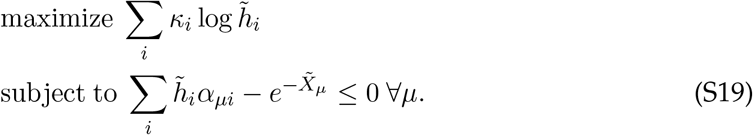

To convince ourselves that these two problems really have the same solution, we return to the equation defining *h*_*i*_ in terms of the relative equilibrium abundances *f*_*µ*_:

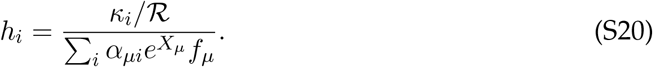

Translating the uptake budgets by 𝒳 rescales *h*_*i*_ by *e*^−𝒳^, while leaving *f*_*µ*_ unchanged. Therefore, both optimization problems correspond to the same *f*_*µ*_, which is the actual observable we are interested in. In our numerical simulations, we solve this optimization problem directly, with an arbitrary fitness gauge chosen for numerical convenience.

Once the *h*_*i*_ have been determined, the strain relative abundances *f*_*µ*_ can be calculated numerically. First, the living strains with *f*_*µ*_ *>* 0 are identified as the strains for which the inequality constraint in Eq. (S17) achieves equality. Given these strains, the definition of *h*_*i*_ in Eq. (S7b) can be inverted to obtain a system of linear equations for *f*_*µ*_:

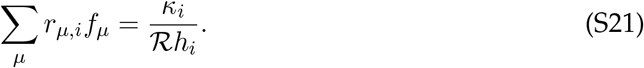

Because *f*_*µ*_ for extinct strains must be zero, there are only 𝒮^*^ ≤ ℓ remaining abundances to be solved for. A non-negative linear least squares solver can be used to find the unique set of *f*_*µ*_ that satisfy all ℓ equations.

#### 1.6 Analytic Approximation of Ecological Equilibrium

By using this gauge symmetry and a small number of additional approximations, we can rewrite this optimization problem in a more analytically tractable form. Indeed, we find that we can put our model in a similar form to that analyzed in Refs. (36) and (37), by choosing a gauge such that ⟨*h*_*i*_⟩ = 1. Let’s define a constant *h*_0_ based on the equilibrium solution in our (arbitrary) original gauge,

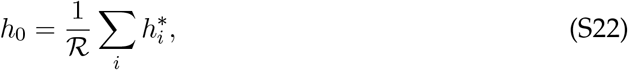

and consider what happens when we set 𝒳 = log *h*_0_. The rescaled solution is

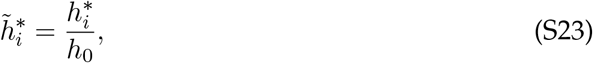

which by definition has mean 1. It is now natural to define a new variable,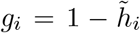, which has mean zero. The optimization problem is now

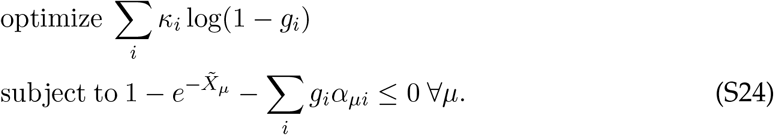

In large ecosystems, the *g*_*i*_’s will often be small compared to one, allowing us to Taylor expand the optimization function to second order. (We will verify the range of parameters where this holds in SI Section 3.5.) When *g*_*i*_ ≲ 1, the third term on the LHS of the constraint equation must also be less than one. This implies that for any surviving species (where the constraint equation is exactly satisfied) the fitness term 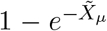 must also be small compared to one. In other words, choosing a gauge where ⟨*h*_*i*_⟩ = 1 is the same as choosing a gauge where the pure fitnesses of surviving organisms are centered at zero, allowing us to Taylor expand the 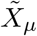 as well. This relationship arises from the fact that strains with very large differences in total uptake budget are unlikely to coexist. Altogether, the approximate optimization problem is

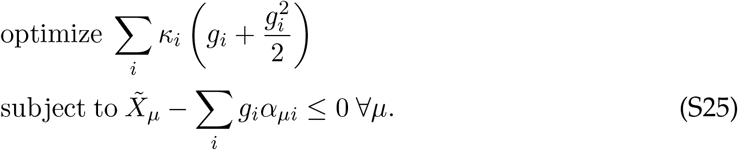

For organisms that go extinct, the Taylor expansion of the 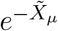 terms may not be very accurate, since 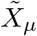 can be arbitrarily negative. However, since 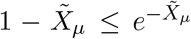, this approximation will not cause us to erroneously label any extinct species as alive – and for the species which are alive, we will check that the approximation is sufficiently accurate to avoid worry. While it is not yet clear why this approximation is useful, we will see below that remapping the consumer resource problem in this way makes it amenable to replica-theoretic analysis. Note that for numerical simulations, we do not make these approximations, instead solving the original optimization problem in Eq. (S17) – as such, our simulation results implicitly test these assumptions across a range of parameters. We discuss the regime of validity of these assumptions and their interpretations in SI Section 3.5.

### 2 Replica-Theoretic Analysis of First-Step Invasion

#### 2.1 Single-Ecosystem Approximation of First-Step Invasion

Having defined our species pool and consumer resource model, we now consider adding first-step mutations into this community. As discussed in the Model section, mutant-parent coexistence can be described as two correlated community assembly processes. Here, we argue that we can reasonably approximate the conditions for mutant-parent coexistence while considering only one community assembly process, which simplifies our theoretical work. First, we describe the two-ecosystem framework. An ecosystem *E*_1_ is formed through standard community assembly: 𝒮 species are placed into competition, and 𝒮^*^ remain alive at equilibrium. Then, one of the surviving species produces a beneficial mutant offspring. This effectively creates a new community assembly problem for ecosystem *E*_2_, where the initial ^*^ species plus one mutant are allowed to reach ecological equilibrium. The mutant-parent coexistence probability can therefore be written as

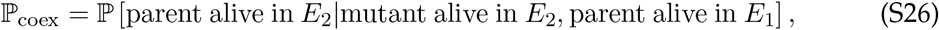

where the conditional probability requires that (1) the mutant be beneficial and (2) the parent be alive when it produces a mutant. Standard manipulation of probabilities allows us to rewrite this as

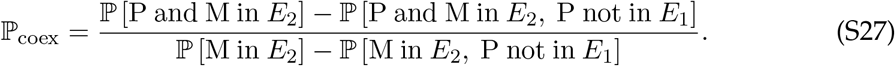

First, we argue that ℙ [P and M in *E*_2_, P not in *E*_1_] ≈ 0. This probability describes a situation where the parent is unable to survive in the initial ecosystem, but its hypothetical mutant offspring is. Moreover, the invasion of the mutant offspring adjusts the ecosystem in a way where the original parent strain is also able to simultaneously invade. Intuitively, both of these requirements are unlikely: it will be difficult for a small mutation to allow an otherwise unfavored species to invade, and we would naively expect that the presence of the mutant would make it more difficult for the parent species to survive – not easier – due to their shared competition for resources. Making this approximation, we find

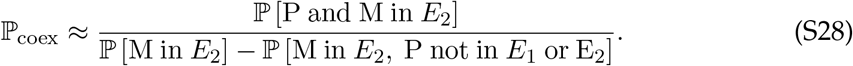

The next approximation we make is similar: we assume that if the mutant is able to survive, the parent would likely be able to survive in its absence. Mathematically, ℙ [M in *E*_2_] ≫ P ℙ [M in *E*_2_, P not in *E*_1_ or E_2_]. While it is possible that a small-effect beneficial mutation could push an organism over the survival threshold, we expect that *most* randomly chosen surviving organisms will not be able to be rescued from extinction by a single mutation. Thus, we write

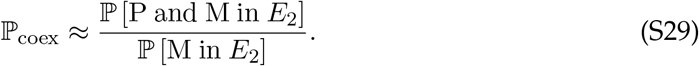

Now, our coexistence probability depends only on one ecosystem. But there is still an issue: *E*_2_ is not a traditionally assembled community, since it is formed only from the survivors of an earlier community assembly process plus a mutant, rather than independently sampled organisms. Let’s instead consider *E*_*s*_, an ecosystem obtained from simultaneous community assembly of 𝒮 independently sampled species, plus one “mutant” species which is closely related to one of the 𝒮 other species. The key difference between *E*_2_ and *E*_*s*_ is that *E*_*s*_ can potentially contain species which were extinct in *E*_1_, but are brought “back to life” by the invasion of the mutant. While such returns are possible (37), we conjecture that the invasion of a closely-related mutant strain should be a relatively small perturbation to the ecosystem, and as such these re-introductions should be rare – particularly when the number of resources and species are large. So, as a first approximation, we do not expect them to meaningfully impact ℙ_coex_. After these approximations, we now have

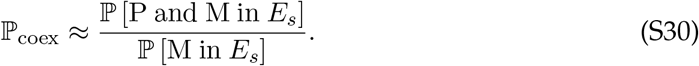

This approximation allows us to estimate the mutant-parent coexistence probability in an assembled community by only considering one community assembly process, which includes two closely correlated organisms. (Which one we have labeled the “parent” and “mutant” is essentially arbitrary after making these approximations.) As shown in Fig. S4, simulations confirm that these approximations are valid across most of the parameter regime we study, so our one-ecosystem approximation produces results which match the two-ecosystem one. An exception is at very high expected niche saturation, where species “coming back to life” after mutant invasion becomes more relevant. However, there is a narrow range of parameters where this effect is significant (particularly for ecosystems with many resources), and our approximation where these species are allowed to return nonetheless sets a lower bound on the mutant-parent coexistence probability.

#### 2.2 Partition Function and the Replica Trick

We would now like to argue that our model can be treated similarly to Refs. (36) and (37), meaning that their results describe a baseline assembled community before we add mutations. While we will not reproduce every element of their calculation, we restate key steps and approximations. Since ecological equilibrium can be described as the solution to an optimization problem, we can write a partition function for a system with energy 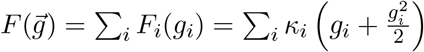 at temperature *β*^−1^. In the zero-temperature limit *β* → ∞, the system will be exactly at ecological equilibrium; at finite values of *β*, fluctuations from equilibrium correspond to shot noise in the population. The partition function is

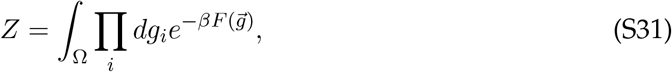

where Ω corresponds to the constrained region in our optimization problem, where resource surplus is non-positive. We can write out this region explicitly using the Heaviside function θ(*x*) = max(0, *x*) as follows:

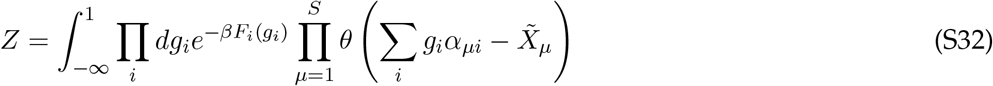

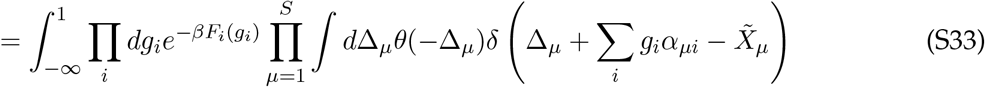

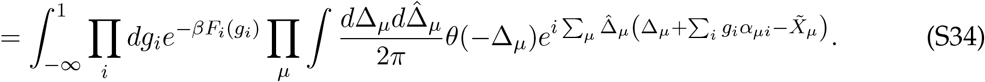

where 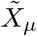 is the pure fitness of species *µ*, in the gauge described in 6.1.2. We have traded our complicated integration region for two additional integration variables for each species: the resource surplus Δ_*µ*_ and its auxiliary Fourier variable 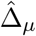.

This partition function is written for a particular choice of *S* sampled species. However, we are interested not in any particular choices of species, but the behavior of the ecosystem when *typical* species are drawn from some random distribution. These typical ecosystems can be analyzed by calculating ⟨ log *Z*⟩, where the average is over our random draws of species. This average can be calculated through use of the replica trick,

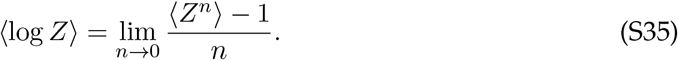

In accordance with the replica trick, we consider *n* copies of our system, which share identical resident species but may differ in shot noise (encoded by the temperature β). Then, we treat *n* as a real number and consider the limit *n* → 0, comparing our results to simulation to ensure this approximation holds. With *n* copies of our system, the average partition function is

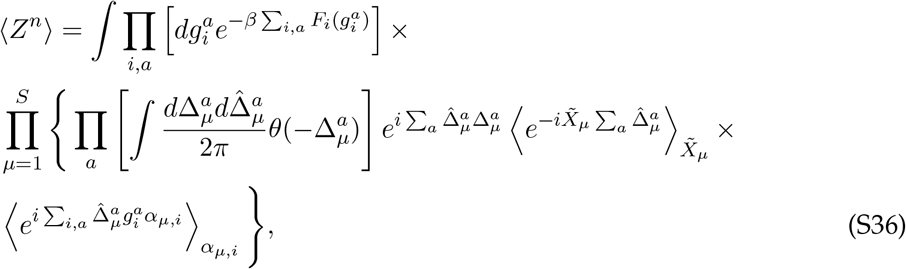

where *a* runs from 1 to *n* and indexes the replicas of our system. Next, let’s average over our random draws of species. Suppose that the pure fitnesses 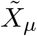 are drawn from a normal distribution with mean 𝒳/*ℛ*_0_ and variance (*ϵ*/*ℛ*_0_)^2^. Recall that 𝒳 represents our fitness gauge, and will later be set to ensure that ⟨*g*_*i*_⟩ = 0. The factors of 1/ *ℛ*_0_ account for the fact that the *α*_*µ,i*_ in Eq. (S25) are ~ 1/ *ℛ*_0_ smaller than the corresponding terms in Refs. (36) and (37) due to the overall normalization of 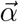. The 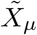-dependent term then becomes

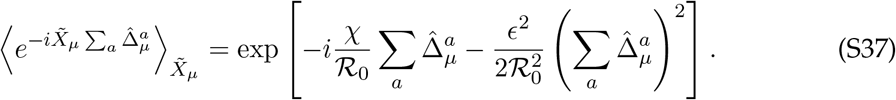

Now, let’s consider the average over the strategy vectors. For binary resource strategies, the *α*_*µi*_ are defined as

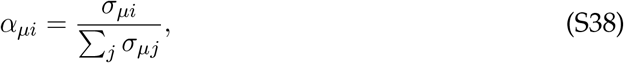

where *σ*_*µi*_ are i.i.d. random variables equal to 1 with probability ℛ_0_/ℛ and 0 otherwise. Unlike the *σ*_*µi*_, the *α*_*µi*_ are not independent for the same *µ* with different *i*’s. However, we can approximate them as independent by separating out the dependent portion, provided that ℛ_0_ ≫ 1:

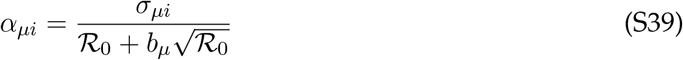

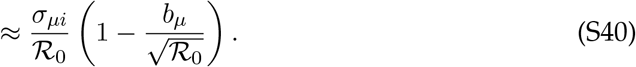

The *b*_*µ*_ are mean-zero 𝒪(1) random variables representing the fluctuations in the total number of resources used by each species. While not technically independent of the *σ*_*µi*_, this dependence would contribute to terms with smaller powers of ℛ_0_ and can thus be ignored. So, we consider their averages separately:

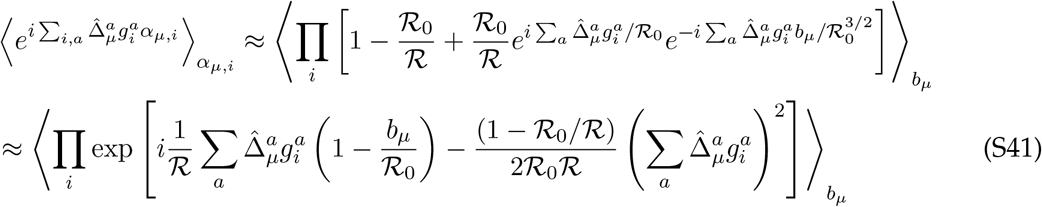

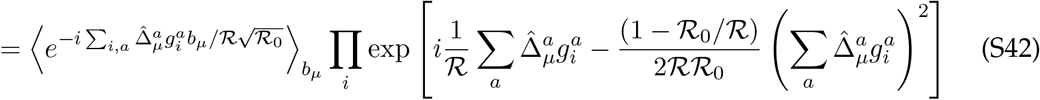

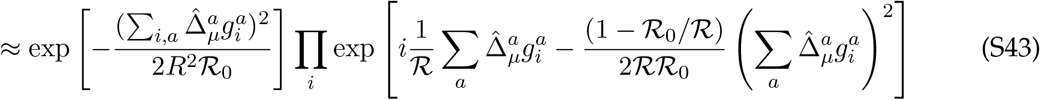

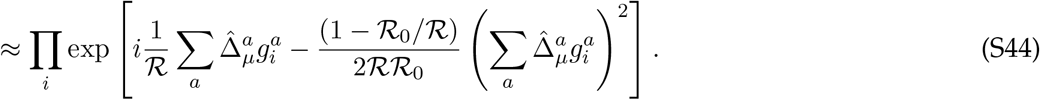

Effectively, this approximation treats 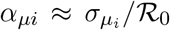. Finally, to bring our answer in line with Refs. (36) and (37), we rescale variables 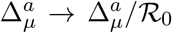 and 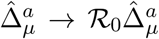. The final partition function is

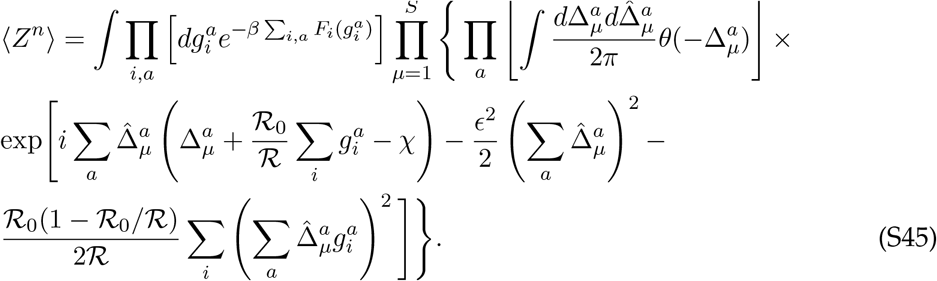

This is the same as the partition function calculated in Refs. (36) and (37), except for the presence of 𝒳. We can ignore 𝒳 until later in the calculation (for example, by absorbing it into the definition of 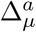, which causes it to appear only within the Heaviside function), and re-introduce it only when we require that ⟨*g*_*i*_⟩ = 0.

#### 2.3 Introducing First-Step Mutations

The above partition function is valid for an ecosystem where all species are sampled randomly. However, in our “simultaneous assembly” approximation of first-step mutation, we would like to include two sampled species which are very related to each other, differing only by a single knock-out mutation. We will index the parent and mutant species *P* and *M* ; they have the same resource consumption strategy, except that the parent uses resource 1, while the mutant’s capacity to use resource 1 is lowered by a factor of γ. For a full knock-out mutation which preserves our assumption about binary resource use, γ = 1, but this more general type of mutation will aid us when interpreting our conclusions later. In particular, we will find that the magnitude of the phenotypic change, 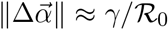, is most important when predicting its impact on the community. We assume the mutant’s pure fitness is increased by a total amount Δ*X* relative to its parent (which can be negative to indicate an overall cost to the mutation). With the parent-mutant pair accounted for, the partition function becomes

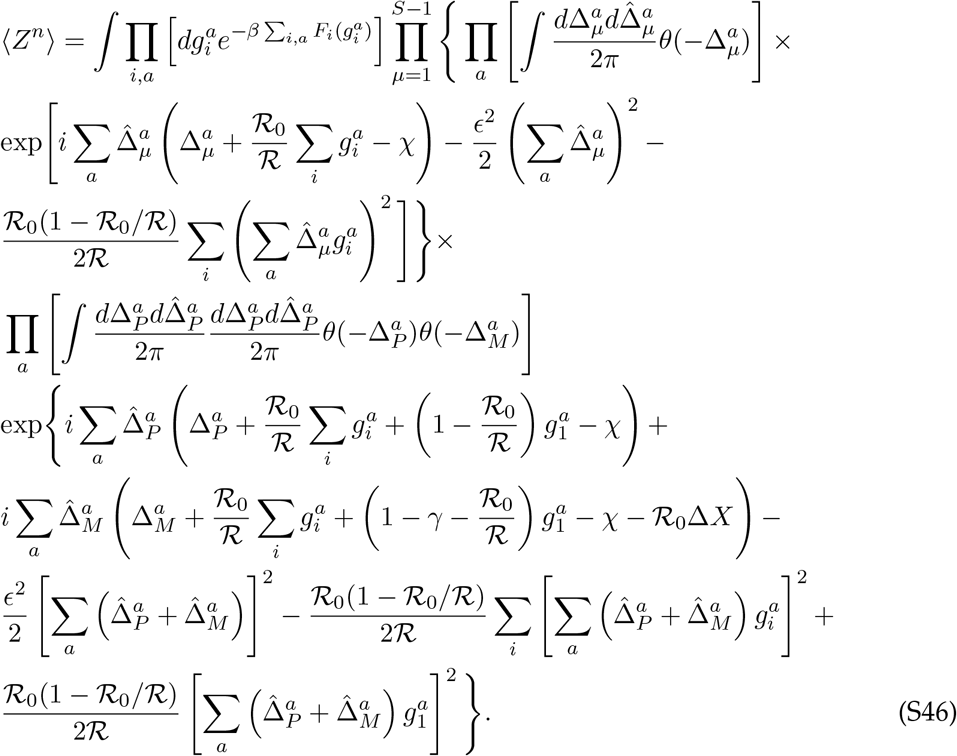

(Note that the Δ*X* term has a factor of ℛ_0_ to account for the rescaling we did earlier.) The next few steps of the calculation are a series of approximations which allow us to evaluate this complicated integral, paralleling those in Refs. (36) and (37).

#### 2.4 Saddle Point Approximations

For ℛ ≫ 1, the central limit theorem dictates that the mean and variance of the equilibrium *g*_*i*_ (which encode the availabilities of each resource) should approach a deterministic value. Introducing the mutant-parent pair is a relatively small change in the ecosystem, so we expect that these deterministic values should be the same as those calculated in previous work, if the mutant-parent pair were not present. This motivates us to define replica-specific “order parameters”

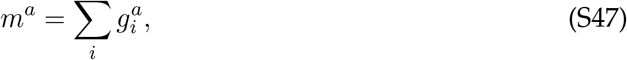

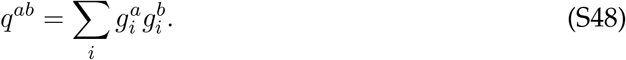

As we did with the resource surplus 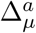, we can introduce these order parameters as new integration variables, defined via delta functions in their Fourier representation. This results in an integral

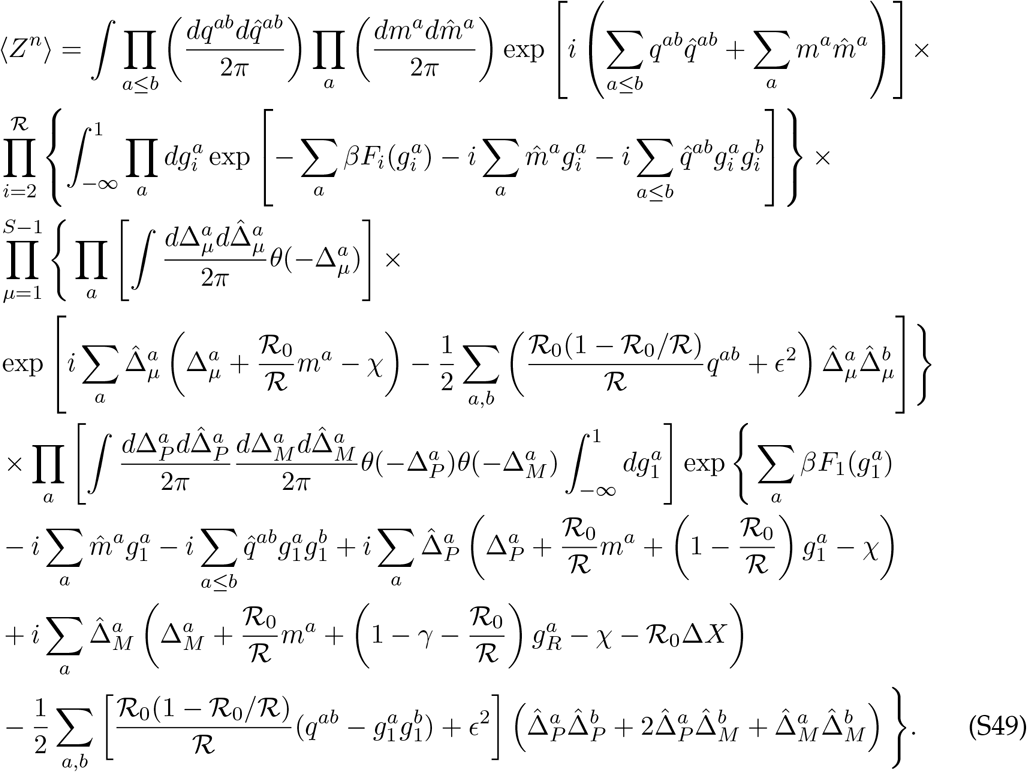

So far, we have made no approximations from our initial partition function. Now, however, we invoke the saddle-point approximation, stating that the probability distribution for the order parameters and their Fourier conjugates is so heavily peaked that their values are equivalent across replicas. In short, we assume

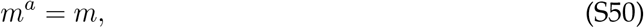

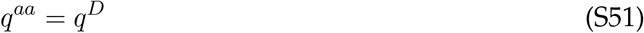

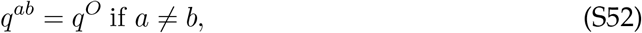

and likewise for the conjugate (hatted) order parameters. We recall that we must eventually choose our fitness gauge *χ* such that *m* = 0, a deviation from the analysis in Refs. (36) and (37) due to our choice of consumer resource model. Our saddle point approximation allows us to significantly simplify our integral, by eliminating the integrals over the replica-specific order parameters. This change decouples our large integral expression into a product of three *independent* integrals: one over the resource availabilities *g*_*i*_ for *i >* 1, one over the surpluses 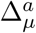 for all species besides the parent and the mutant, and the final integral over *g*_1_ and the surpluses for the mutant and parent strains.

Because we assume a diverse, many-species ecosystem, the species besides the parent and mutant will be most important in setting the saddle-point values of our order parameters and their Fourier conjugates. Thus, we approximate that these saddle-point values are unchanged from the calculation in Refs. (36) and (37), which includes only the first two sets of integrals (i.e., those independent of the parent and mutant). While we will not restate their calculation here, we will summarize their results. If we assume that the values of *q*^*D*^ and *q*^*O*^ are similar –

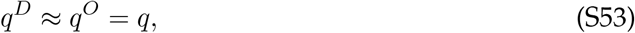

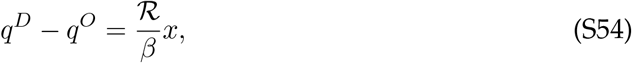

then the saddle-point values of the conjugate variables are

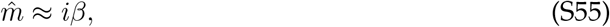

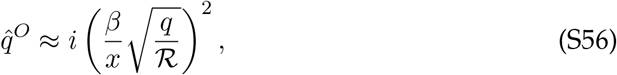

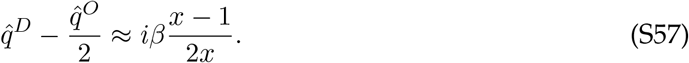

These equations leave *q* and *x* undetermined, which can be solved for implicitly and approximated through numerical equations which we will discuss later.

We also make the approximation 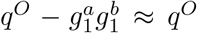. The order parameter *q* is formed by adding together 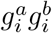 for each of ℛ resources, and we expect each of these quantities to be positive. So, the scale of 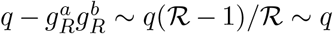. After making this approximation, let us also recall that 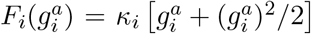. Knowing that *m* = 0 and 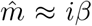 causes all the terms linearly dependent on the 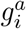 to cancel out, besides the mutation-dependent 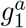 terms: thus, before considering the effects of the mutation, our system is invariant under rotation of the 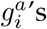. Practically, this means that our initial choice of a knock-out mutation was arbitrary: a knock-in mutation, or any strategy change 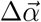 with the same overall magnitude, can be redefined as a change in a single element through an appropriate rotation. Our results should therefore be valid for mutations which affect multiple resources simultaneously.

#### 2.5 Joint Distribution of Δ_*P*_ and Δ_*M*_

Now that we have decoupled our integrals through the saddle-point approximation, we are nearly ready to analyze the phenomenon of parent-mutant coexistence. To do this, we only care about the part of the partition function which depends on those strains. For simplicity, we will also at this point assume uniform resource supply (*κ*_*i*_ = 1); we will discuss at the end of our calculation what happens when this assumption is relaxed. With the saddle-point approximation for *κ*_*i*_ = 1, the portion of the integral in Eq. (S49) which depends on the parent and mutant strains is

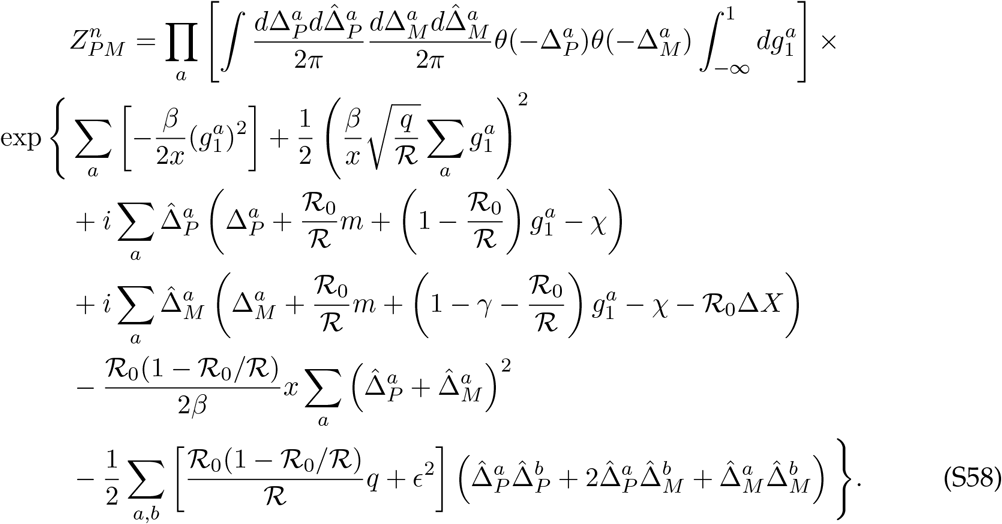

Next, we perform a change of variables to 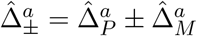:

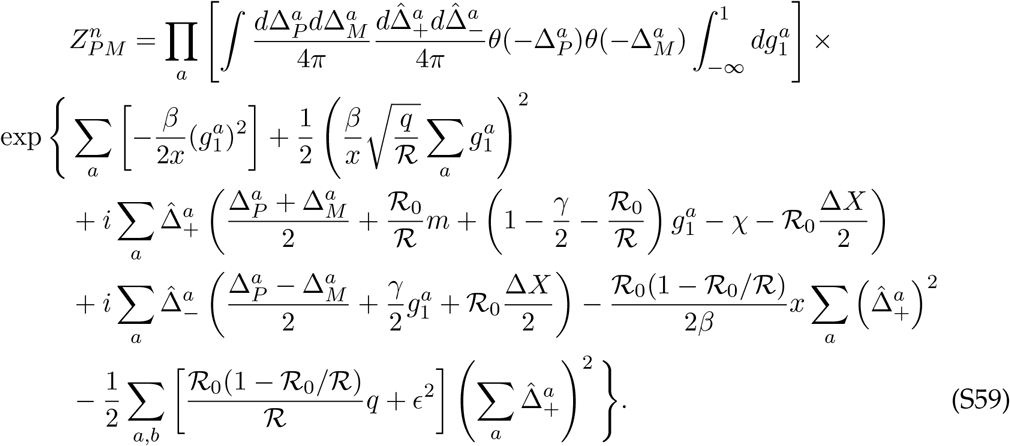

We now aim to decouple the replicas, by eliminating terms which involve more than one replica. All these terms are of the form (∑_*a*_ *f*^*a*^)^2^, where *f*^*a*^ is some quantity which depends only on one replica. We can decouple these replicas using the identity

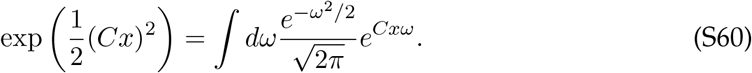

Thus, we can decouple the replicas at the cost of introducing an additional Gaussian integration variable per square term. We can now drop the *a* indices and replace them with a simple power to *n*:

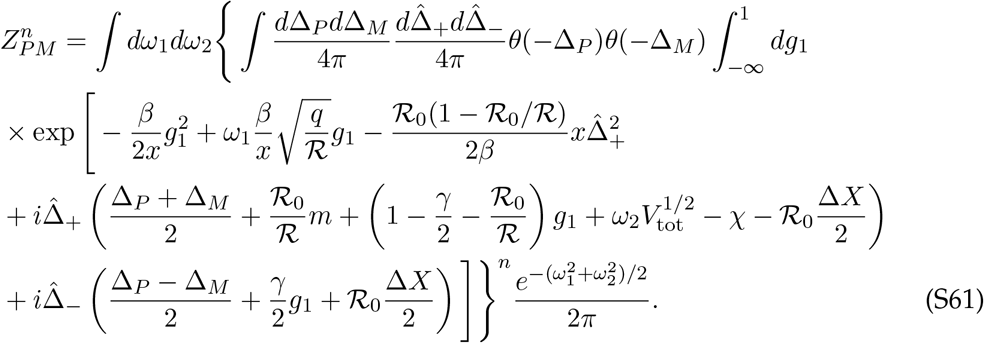

Here, we have defined *V*_tot_ = *q ℛ*_0_(1 ℛ_0_/ℛ)/*ℛ*+ *ϵ*^2^. This parameter, defined as *ψ*^2^ in Refs. (36) and (37), represents the total variance in fitness among sampled species from both strategy and pure fitness. As we will demonstrate later, the integration variables *ω*_1_ and *ω*_2_ do in fact have physical meaning: *ω*_1_ is related to the invasion fitness of the mutant, while *ω*_2_ is related to the abundance of the parent. Next, we perform the integral over *g*_1_. To do so, we assume that most of the weight of the integral is concentrated near small *g*, meaning that we can approximate the integration bounds as −∞ to ∞ rather than −∞ to 1. We also condense all of the resulting terms which do not depend on 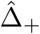 or 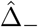 into a constant term *c*.

We can write the result as

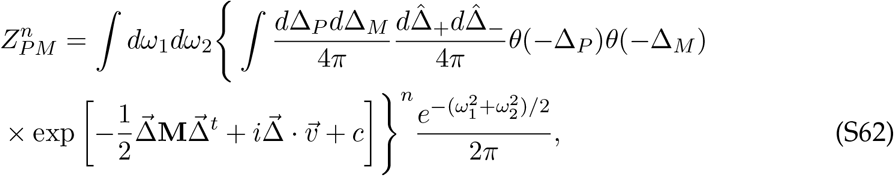

where

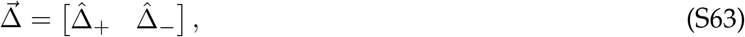

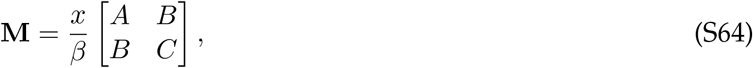

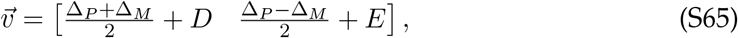

and we define

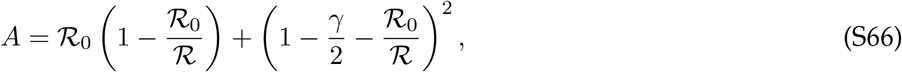

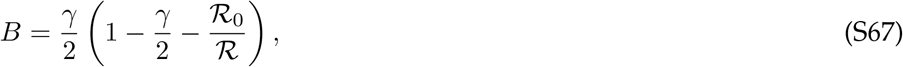

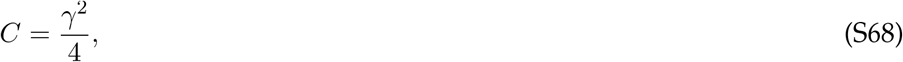

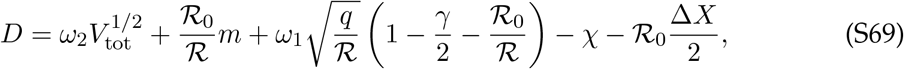

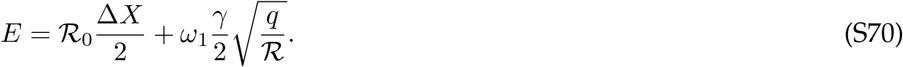

We can now perform the integrals over 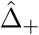 and 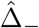. This yields

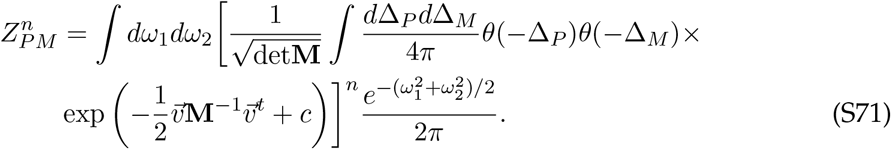

Now, let’s utilize the replica trick limit *n* → 0. Again following the methodology in Refs. (36) and (37), this can be used to recover the joint probability distribution *ρ*(Δ_*P*_, Δ_*M*_):

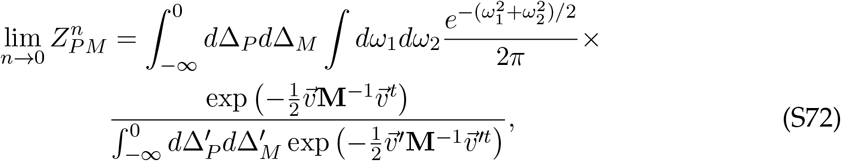

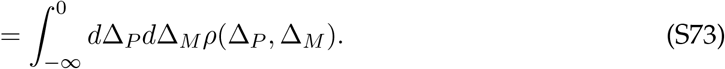

Recall that **M**^−1^ ∼β, an inverse temperature which scales with the number of cells in our community, reflecting the shot noise in our population at any given time. If we assume that our population is always exactly at equilibrium (i.e., is large enough that shot noise is negligible), we can take the *β* → ∞ limit, in which case the exponentials involving **M**^−1^ will be very strongly peaked. In this limit, we can therefore approximate the ratio in Eq. (S72) as a delta function:

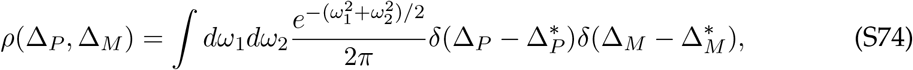

where 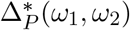 and 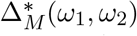 are the non-positive values of Δ_*P*_ and Δ_*M*_ that minimize 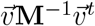 for a given *ω*_1_ and *ω*_2_.

#### 2.6 Parent-Mutant Coexistence Probability

The parent-mutant coexistence probability is encoded in the joint distribution *ρ*(Δ_*P*_, Δ_*M*_):

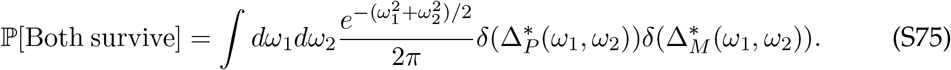

So, the next step is to uncover the functional forms of 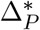 and 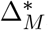, which represent the values of Δ_*P*_ and Δ_*M*_ that minimize 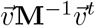, constrained such that both Δ_*P*_ and Δ_*M*_ are non-positive. The matrix **M** and its inverse are positive definite, so this is a convex optimization problem, and either the unconstrained minimum lies in the interior of the allowed region, or the function is optimized on the boundary. If the global minimum happens to be within the allowed region, corresponding to both the parent and the mutant being extinct, these quantities are simple:

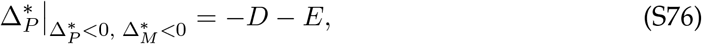

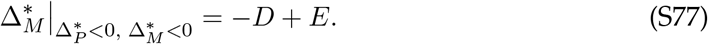

This unconstrained minimum will be in the allowed region if both of these quantities are negative, which occurs when *D >* |*E* |. We see that when both the parent and mutant are extinct, *D* tells us the resource deficit averaged between the parent and mutant, while 2*E* is the difference in resource deficit between mutant and parent. If the average resource deficit is too large, then neither strain can survive, thus

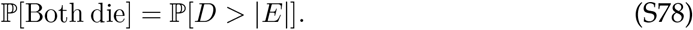

If the unconstrained minimum lies outside the allowed region, then the function will be minimized on the boundary of the allowed region, when one or both of the parent and mutant are alive. For example, if the mutant survives and the parent goes extinct, the minimum will be found at 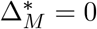 and

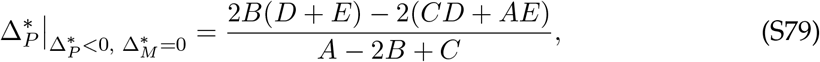

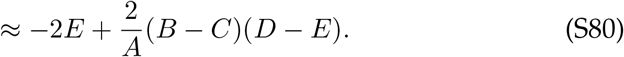

This expansion comes from the fact that *A* ∼ ℛ_0_ (provided that ℛ_0_ does not get too close to ℛ), while *B, C* ∼ 1. Similarly, if the mutant goes extinct while the parent survives, we find

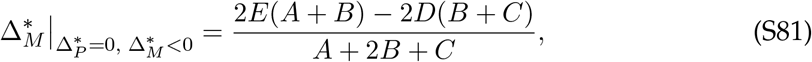

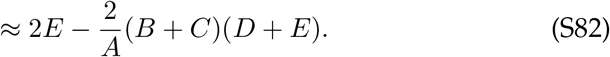

The last possibility is that the optimum lies on the corner of the allowed region, where the parent and mutant coexist (so 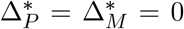). We can find this probability by requiring that each of these three potential optima are outside of the allowed region, which occurs when

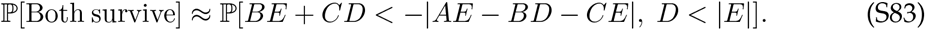

Recalling that *A* ∼ ℛ_0_ ≫ 1, the *AE* term dominates unless |*E*| ≪ 1, meaning that *E* must be small for the inequality to be possible. (Note that the ℛ_0_Δ*X* term is not actually 𝒪 (ℛ_0_), because the natural fitness scale for a mutant turns out to be 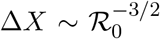) When this is the case, the *BE* and *CE* terms will be even smaller, allowing us to safely ignore them:

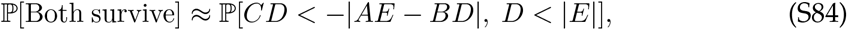

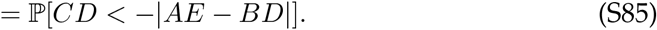

We have eliminated the second condition because it is now redundant with the first: since *C >* 0, the first inequality requires *D <* 0, trivially satisfying *D <* |*E*|. Finally, let’s recall that in our simultaneous assembly approximation of mutant invasion (Appendix 1.3), we must condition on the probability that the mutant survives:

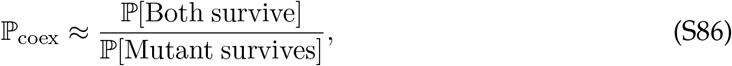

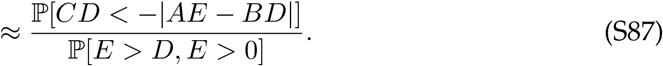

In principle, this is a quantity that can be calculated numerically. However, this result is difficult to intuitively interpret in this form, and it still depends on the saddle-point order parameter *q*. Nonetheless, we can immediately make two meaningful conclusions. If we send the number of resources consumed by a typical organism ℛ_0_ → ∞, then *A* becomes large and ℙ_coex_ → 0. Likewise, if we send the effect size of our mutation *γ* → 0, then 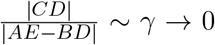, and thus ℙ_coex_ → 0. Since we are considering a mutation whose effect size is comparable to changing one resource’s uptake rate by a factor of γ, both of these observations suggest the same thing: if the phenotypic effect of a mutation is too small, mutant-parent coexistence is impossible. This phenomenon is a signature of “mesoscopic” mutation, suggesting that our results may not be captured by models of mutations with infinitesimal phenotypic effects, such as adaptive dynamics approaches (44, 65).

### 3 Scaling Analysis of First-Step Mutations

In this section, we aim to restore physical intuition to our replica-theoretic results, finding approximate expressions which indicate how the frequency of inter-niche evolution scales with community parameters such as the number of surviving species.

#### 3.1 Sampling Depth and Fitness Gauge

We must utilize the saddle-point results for an assembled community without mutation, largely from Refs. (36) and (37), to calculate our order parameters *m* and *q*. However, we opt for a different set of independent variables, defining our assembly process in terms of the number of sampled species 𝒮 and the typical number of surviving species 𝒮^*^. The community assembly parameter *ϵ*, which describes the variation in pure fitness among sampled species, will then be defined implicitly as a function of these parameters. This change of variables is helpful because 𝒮^*^ is a measurable property of an experimental ecosystem, while *ϵ* describes a property of *sampled* species, which may be very different from typical surviving species and is thus more difficult to access experimentally.

With this set of variables, a natural ratio to consider is the “sampling permissivity” 𝒮^*^/𝒮, roughly the probability that a new independently sampled species would be able to invade the ecosystem. It turns out that it will be useful to define an effective “sampling depth” λ, which is a function of 𝒮^*^/𝒮:

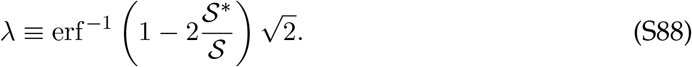

For an undersampled system (𝒮^*^/𝒮 → 1), the sampling depth *λ→* − ∞ ; for an oversampled system (𝒮^*^/𝒮 → 0), we have *λ→* ∞. From the saddle-point results of Refs. (36) and (37) (which we do not re-derive here), we know that

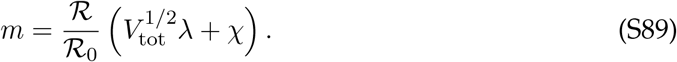

Recall that 𝒳 sets our choice of fitness gauge: the uptake budget of sampled species is drawn with mean 𝒳/ℛ_0_ and standard deviation *ϵ*/*ℛ*_0_. But in our community assembly approximations, we have chosen our gauge such that ⟨*g*_*i*_⟩ = *m*/*ℛ* = 0. This implies that

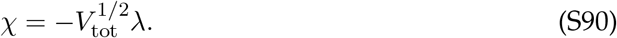

As we sample more species and *λ* increases, the uptake budgets or pure fitnesses of surviving species will be further and further into the high-fitness tail of the distribution. Since the typical pure fitness of surviving species is nearly zero in our fitness gauge, *χ* must become more negative as *λ* increases to compensate for the increased sampling. In the consumer resource model analyzed in Refs. (36) and (37) (where *χ* = 0), an increase in sampling depth results in an overall increase in *m*: as oversampled species become higher and higher in pure fitness, the typical resource becomes less valuable to consume. This effect causes resource knock-out mutations to be more beneficial on average than knock-in mutations in the oversampled limit. In our model, we adjust the “zero” of pure fitness to compensate for this effect, such that *m* = 0 regardless of *λ*.

For future sections, it is helpful to define a function *I*(*λ*) such that

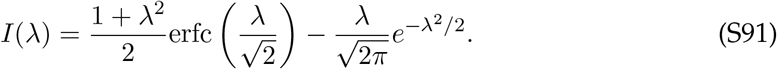

#### 3.2 Niche Saturation and *σ*_inv_

From saddle-point results in Refs. (36) and (37), we can write expressions for *V*_tot_, *q*, and *ϵ* in terms of quantities we have already defined, as well as variance in resource supply

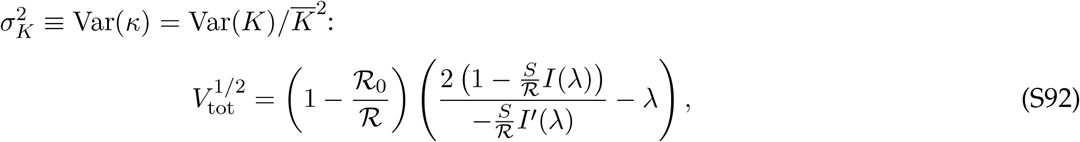

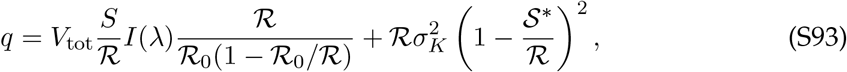

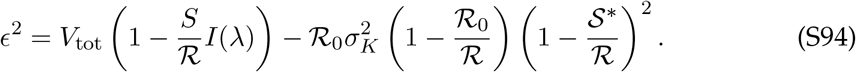

The order parameter *q* describes the total variance in resource availability, allowing us to find the distribution of the resource availabilities *h*_*i*_: a Gaussian with mean 1 and variance *q*/ℛ. A mutation changing use of a single resource by a factor γ will have an invasion fitness with magnitude γ|1 − *h*_*i*_|/ℛ_0_. This leads us to define the quantity

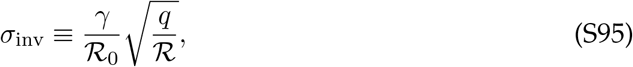

which describes the typical magnitude of the invasion fitness of a strategy mutant 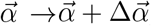, where 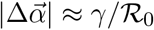. (For example, for a complete knock-out mutant for a system with binary resource use, we have *γ* = 1.)

While these equations completely determine all the variables we have defined, they are difficult to interpret physically, having complicated dependence on the ratios 𝒮/ℛ and ℛ_0_/*ℛ* as well as the sampling depth *λ*. We can make progress by considering how these equations depend on the *niche saturation* 𝒮^*^/ℛ ≤ 1, describing how close to maximum capacity we expect our final ecosystem to be. By rewriting 𝒮/ℛ = (𝒮/𝒮^*^)(𝒮^*^/ℛ), we separate out the parts of these equations that depend on the sampling depth **λ** (or equivalently, 𝒮^*^/𝒮) from the niche saturation 𝒮^*^/ℛ. (Technically, rather than using the value of 𝒮^*^ for any particular ecosystem as an input parameter, we use its expectation ⟨𝒮^*^⟩ over random ecosystems with the same underlying parameters. Since these values tend to be close in practice, as shown in Figure 2B, we will simply write 𝒮^*^ in our scaling relationships.)

To simplify our results, it is helpful to expand our expressions in the undersampled and oversampled limits (*λ*→ ± ∞). It is worth noting that the *λ*→ − ∞ limit for fixed 𝒮^*^/ℛ does not correspond to a physical community assembly process: the expression for *ϵ* in Eqn. (S94) must be positive, but (𝒮/ℛ) ·*I*(*λ*) → ∞ as *λ*→ − ∞, making the RHS of that equation negative. Effectively, this means that in order to sample species such that a desired fraction of niches are filled, a minimum fraction of sampled species must go extinct in the final ecosystem (on average). However, many quantities we are interested in are still well-defined in the *λ*→ −∞ limit, and indeed scale similarly to the *λ*→ ∞ limit: we use this to demonstrate an overall lack of dependence on sampling depth.

We first quote results for 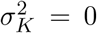 From standard expansions of the error function, we obtain

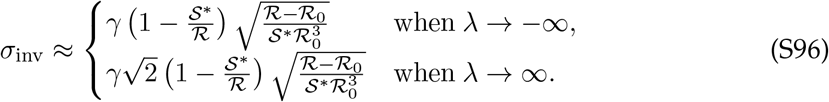

Remarkably, the typical invasion fitness scale of a strategy mutation has very weak dependence on the sampling depth *λ*, changing only by a factor of 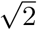 from the extremely undersampled to the extremely oversampled regimes. This result allows us to effectively drop out *λ* (and thus 𝒮^*^/*𝒮*) as a variable, which was very difficult to see from our saddle-point results above.

#### 3.3 Scaling Analysis of ℙ_coex_

We got our expression for *σ*_inv_ “for free” from the saddle-point value of *q*, without need to explicitly consider the impact of the parent and mutant on the ecosystem. The same is not true for the mutant-parent coexistence probability: to obtain an interpretable expression for ℙ_coex_, we must start from our results in Appendix 2.5, and then undergo a similar procedure of plugging in saddle-point values in the *λ*→ ± ∞ limits. Recall that in our two-ecosystem approximation, we have

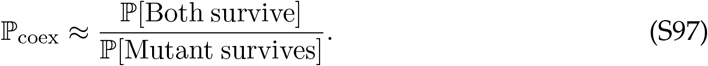

To get some intuition, let’s start with the simpler calculation of the denominator of this expression: the probability that the mutant species survives in our assembled ecosystem. From our earlier results, this is

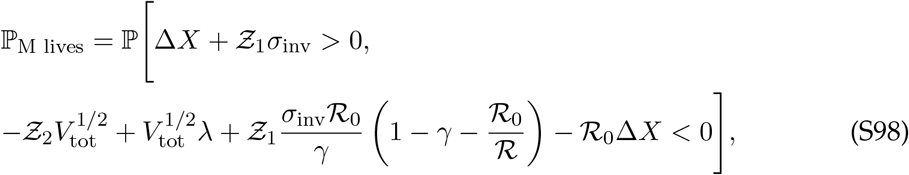

where *Ƶ*_*i*_ are independent standard normal random variables. Note that we have added a negative sign to *Ƶ*_2_; while this leaves its distribution unchanged, it will aid in interpretation. The first two terms in the second inequality scale with the fitness variation in all sampled species 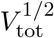, while the remaining terms scale with the fitness change induced by the mutation. If the mutation results in a small change in the organism’s overall fitness, the first two terms should dominate, so we can approximate

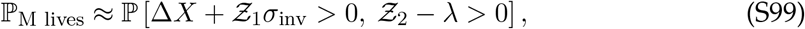

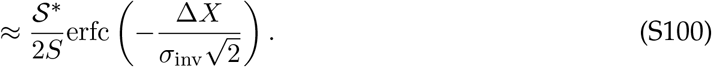

Each of these inequalities has a meaning which allows us to interpret *Ƶ*_1,2_ as physical quantities. The first inequality represents the probability the mutant is beneficial relative to the parent: the chances that the deterministic fitness change Δ*X* combined with the stochastic fitness change *Ƶ*_1_*σ*_inv_ due to the strategy mutation is positive. The second inequality represents the requirement that the parent background the mutant arises from is well-adapted enough to survive in the population in the first place: only draws of *Ƶ*_2_ which are greater than *λ* will survive in the population. Indeed, the probability that *Ƶ*_2_ *> λ* is simply 𝒮^*^/𝒮, the probability a randomly chosen species would survive in the community. Neglecting the other terms in the second inequality neglects the possibility that the parent cannot survive in the initial population, but the mutant can; as discussed in SI Section 2.1, we expect this outcome to be rare.

We posit that *Ƶ*_2_ determines not only the survival of the parent species, but also its approximate relative abundance *f*_*P*_. Past research (29) suggests that consumer resource models of this form yield a truncated Gaussian distribution of abundances, which (up to an overall normalization factor) coincides with the distribution of *Ƶ*_2_ conditioned on being greater than *λ*. We can find this normalization factor by enforcing ⟨*f*_*P*_ *∣f*_*P*_ *>* 0⟩ = 1/𝒮^*^, which yields

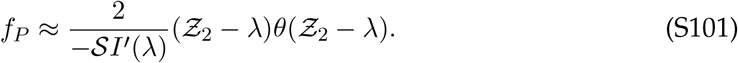

In the oversampled limit *λ*≫ 1, the prefactor is approximately *λ*/𝒮^*^. Our results in the next section confirm that using this relationship between *Ƶ*_2_ and *f*_*P*_ yields reasonably accurate predictions.

Now, let’s calculate the numerator of ℙ_coex_, the probability that the mutant and parent will both be alive in our simultaneously assembled ecosystem. Again approximating that terms which scale with overall fitness variation 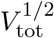 dominate terms which scale with mutant fitness change Δ*X* and*σ*_inv_, we find that

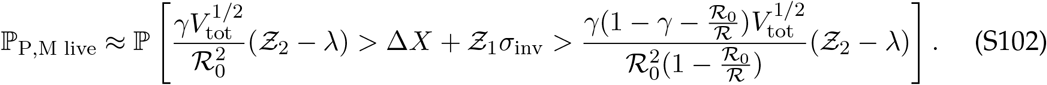

Let us analyze this probability using our physical interpretations of *Ƶ*_1_ and *Ƶ*_2_. The outer inequality only holds when *Ƶ*_2_ *> λ*, again requiring that the parent species survives. The middle term is the leading contribution to the invasion fitness of the mutant, which is 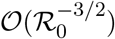. However, the inequality bounds it between two quantities of size 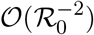. So, we conclude that parent-mutant coexistence is achieved when the invasion fitness of the mutant is sufficiently small.

Since the allowed region for *Ƶ*_1_ is smaller than its typical width by a factor of 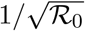, we can approximate its probability density as constant over the allowed region. This gives us

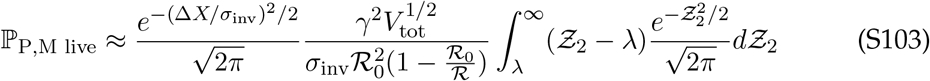

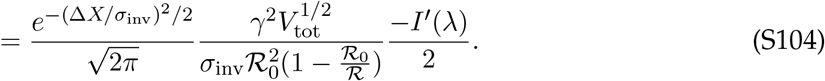

Finally, we divide by the probability the mutant lives, and consider the undersampled and oversampled limits when Δ*X* = 0:

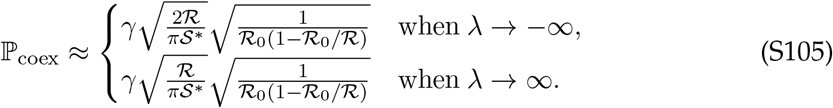

Once again, the dependence on the sampling depth is nearly negligible, contributing only an 𝒪(1) numerical factor. Note, however, that this result is different from that in the main text in Equation (9). This is because in our theoretical calculation, we arbitrarily chose the identity of the parent species and the mutation, effectively giving each surviving species and each mutation an equal chance of being selected. In reality, each surviving species produces mutations at a rate proportional to its abundance, and mutations have a probability proportional to their invasion fitness of surviving genetic drift (52). In the next section, we will discuss how to correct for this difference.

#### 3.4 Dependence of ℙ_coex_ on Invasion Fitness and Abundance

Let us reframe our expression for ℙ_coex_ in terms of the abundance *f*_*P*_ of the parent and the invasion fitness *s*_inv_ of the mutant, relating them to *Ƶ*_1_ and *Ƶ*_2_ as we did earlier. We can read off the invasion fitness of a beneficial mutant as the saddle-point value of 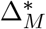 when Δ_*P*_ is fixed at zero, representing the initial growth rate of the mutant *before* it has a chance to invade:

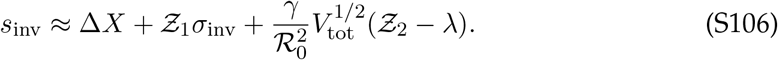

We can plug this expression into our earlier probability that both the mutant and parent live to find that

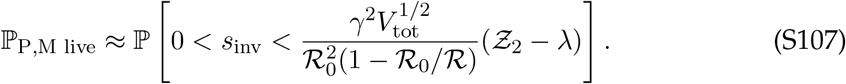

Recall that *Ƶ*_2_ −*λ* is proportional to the abundance of the parent, as we calculated in Eq. (S101). While *Ƶ*_2_ is not independent of *s*_inv_, its contribution to *s*_inv_ is small, so approximating them as independent is reasonable. Plugging in this relationship, we find that

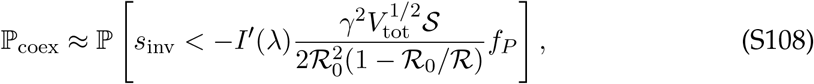

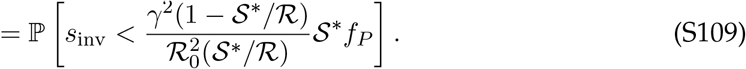

So, our analysis of mutant-parent coexistence has an easily-interpreted meaning: a mutant strain will coexist with its parent when its (positive) invasion fitness is below

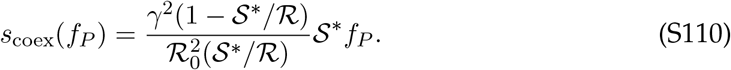

We can calculate this probability by treating *s*_inv_ and *f*_*P*_ as independent random variables, accounting for the probability the mutant survives genetic drift. In this case, their probability densities scale as

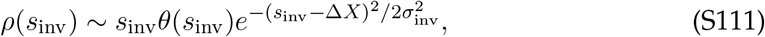

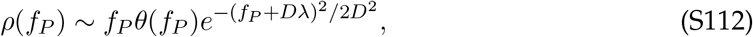

where *D* ≡ − 2/*𝒮I*^*′′*^(*λ*). Finally, ℙ_coex_ can be calculated using standard analytic integration methods, giving the results quoted in the main text after applying similar scaling analysis to the previous section. We can also fix either *s*_inv_ or *f*_*P*_ to calculate a conditional coexistence probability. For example, plugging in a “typical” *f*_*P*_ = 1/𝒮^*^ indicates that 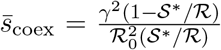 is the typical value of *s*_inv_ above which the coexistence probability begins to fall.

#### 3.5 Self-Consistency of Replica Approximations

Now that we have expressed the results of our theory in terms of physically interpretable parameters, we can use them to check when the assumptions underlying our analytic approximations are valid. The central assumption we made in SI Section 1.5 was that the resource availabilities *h*_*i*_ are not too far from 1, which should fail at some point in the vicinity of |*g*_*i*_| = |1 − *h*_*i*_| ≲ 1. We calculated the typical spread of *g*_*i*_ in terms of *σ*_inv_ in SI Section 3.2, and found that

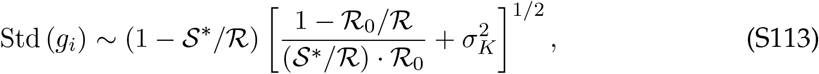

up to an 𝒪(1) factor dependent on 𝒮^*^/𝒮. The first term in the brackets should be small over a wide range of parameter values, since ℛ_0_ ≫ 1. Notably, however, this term causes the |*g*_*i*_| ≲ 1 approximation to break down when the niche saturation is too small, around

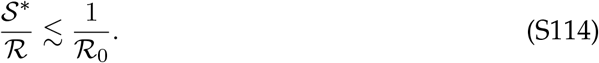

When this inequality is approached or satisfied, there will not be enough species in the community to cover the space of available resources (ℛ_0_ 𝒮 ^*^ ≲ ℛ), so the availability of each resource will be determined by 𝒪 (1) rather than many species. Our simulations provide a test of how approaching this bound affects the numerical accuracy of our results: for example, the leftmost points in Fig. 3B (corresponding to ℛ_0_ 𝒮 ^*^ = ℛ 4) show that the mutant-parent coexistence probability increases more slowly than expected as 𝒮 ^*^/ℛ declines. Similar quantitative deviations from theory occur in more saturated communities when ℛ_0_ is relatively small (e.g. the open circles in Fig. 2D), consistent with our scaling prediction. However, these deviations do not affect our qualitative conclusion that mutant-parent coexistence declines weakly with niche saturation over a wide range of community sizes. Rather, they indicate that in communities where each resource is exploited by only a small number of species, alternative theoretical approaches should be employed.

Self-consistency also requires *σ*_*K*_ ≲ 1, so that the spread in resource supply is not too large. We find that our results hold reasonably well even for *σ*_*K*_ = 1 (Fig. S7B-C). However, different approximations must be employed when the abiotic environment has extremely large asymmetries in resource supply. One such example could be some models of crossfeeding, where an 𝒪 (1) number of externally supplied metabolites result in the production of many other resources at much lower rates.

In our model, a consequence of small |*g*_*i*_| is that the total uptake budgets of surviving species have a maximum spread, Std (*X*_*µ*_ |*f*_*µ*_ *>* 0) ≲ 1, which justified the Taylor expansion in Eq. (S24). In other words, species whose uptake budget differences would drive large changes in their relative abundance over a single generation are unlikely to coexist in the community steady-state. Although this condition places bounds on the spread of *X*_*µ*_ among surviving species, it does not imply that the dynamics are effectively neutral (as in Ref. 32). True neutrality requires that fitness differences are small compared to inverse population sizes and inverse equilibration times; because we focus on large populations and long times, there is a large gap in scales between *X*_*µ*_ ≲ 1 and the emergence of effective neutrality.

The existence of this gap implies that even modest differences in the total uptake budget can still have a significant impact on community structure. Recall that in the simplest version of our model, the uptake budgets of sampled species are drawn from a normal distribution with standard deviation Std(*X*_*µ*_) = *ϵ*/ℛ_0_. The total scaled fitness variation *V*_tot_ among sampled species comes from this variance in uptake budget along with resource consumption strategies. If the ratio *ϵ*^2^/*V*_tot_ ≈ 1, the fates of species will be strongly dependent on their uptake budgets; if *ϵ*^2^/*V*_tot_ ≪ 1, species fates will mostly depend on their specific resource consumption strategies. From the replica-theoretic results in SI Section 3.2, for uniform resource supply, we have

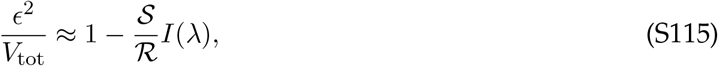

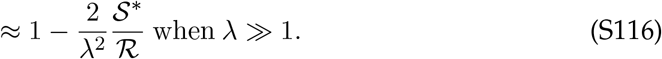

So, when enough species are sampled (large *λ*), total uptake budget will be an important determinant of species fate for any fixed value of 𝒮^*^/ ℛ. Note that if the spread in pure fitness among sampled species is taken to be very small (*ϵ* ≪ 1), then differences in total uptake budget are negligible until enough species have been added to the community to saturate it (𝒮^*^ ≈ ℛ), as in the top curve in Fig. 2B (36, 37). However, our replica-theoretic assumptions continue to hold for *ϵ >* 1, because *surviving* species have a narrower spread of *X*_*µ*_ than *sampled* species:

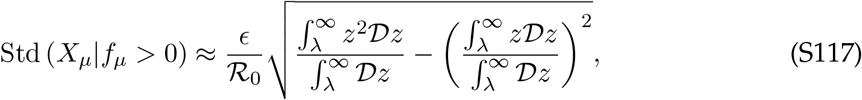

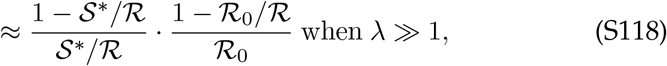

for uniform resource supply, where 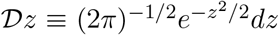 is the standard Gaussian integration element. As expected, the spread in fitnesses becomes large when the spread in the *g*_*i*_’s does, around 𝒮^*^/ ℛ ≲ 1/ℛ_0_, at which point the replica theory fails due to the number of species alive being too small for a “large-ecosystem” approximation.

#### 4 Extensions of Model Assumptions

In this section, we test whether our qualitative conclusions strongly depend on the underlying assumptions of our model, such as the distribution of resource supplies and the method of species sampling. We find that many such changes can be captured by straight-forward extensions of our theory, or simply by using the result from the binary consumer resource model directly.

#### 4.1 Non-Uniform Resource Supply

First, let’s consider non-uniform resource supply (nonzero 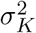), as in (37). If each resource has a supply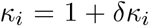, then the availability *h*_*i*_ of that resource will have a component which scales with 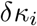 in addition to a community-dependent component. Thus, mutations that knock out a highly-supplied resource will be less beneficial on average. We can determine the strength of this effect by starting with the integrals over the 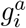 in the second line of Eq. (S49), where the resource supply enters through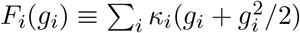. This portion of the partition function was labeled *A*_*i*_ by Ref. (37) (see their Appendix 2, section 1.3). We can follow their calculation to obtain

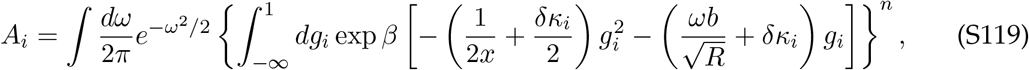

where *b* is a term independent of *δκ*_*i*_ whose value does not concern us here. The contents of the brackets are a Gaussian integral over the *g*_*i*_, centered where the integrand attains its maximum value. Taking non-uniform resource supply, *δκ*_*i*_ ≠ 0, will shift this saddle-point value of *g*_*i*_ by approximately *xδκ*_*i*_ ≪ 1. The saddle-point value of *x* was calculated in Ref. (36) as *x* = 1 𝒮^*^/ ℛ, following our typical substitution of (𝒮^*^/ ℛ) (𝒮/ 𝒮^*^) for 𝒮/ ℛ. This shift in the central value of *g*_*i*_ will shift the invasion fitness of a knock-out mutation targeting that resource by a corresponding amount, since the invasion fitness of a knock-out mutation is simply *g*_*i*_/ℛ_0_. Thus, we can write

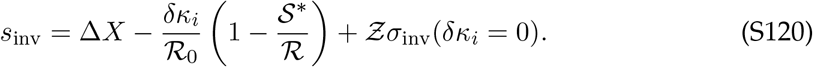

for a mutation knocking out resource *i*. The first term represents the mutation’s change in pure fitness, the second term reflects that environmental supply makes some mutations more or less beneficial on average, while the stochastic third term depends on the specific draws of species in the community, as we calculated earlier. We can quantify the relative importance of environment and community by considering the invasion fitness of a mutation in a monoculture community, where invasion fitness *only* depends on the environment:

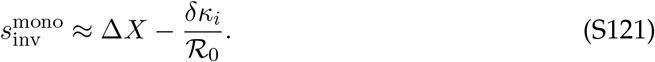

The correlation between the invasion fitness of a mutant in monoculture and in the community context illustrates how much the impact of environmental supply is felt through the “shielding” effects of the community: in the oversampled limit (*λ* → ∞), we find that

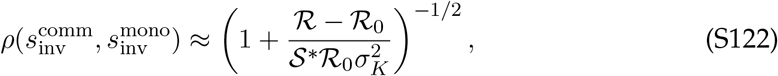

assuming that *δκ*_*i*_ are drawn from a distribution with mean zero and variance 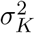. In general, non-uniform resource supply will increase the variance in invasion fitness *σ*_inv_:

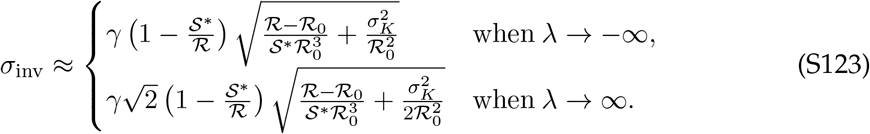

Note that *σ*_inv_ still scales with an overall factor of 1 − 𝒮^*^/ℛ, leaving our previous qualitative results unaffected. Increasing resource supply variation increases *σ*_inv_ and thus lowers the coexistence probability, by making it less likely that the invasion fitness of an arbitrary beneficial mutation will be small enough to make coexistence possible. Although the the-oretical approach assumes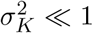, its quantitative predictions hold reasonably well even When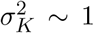, as in the case of exponentially distributed *K*_*i*_ (Fig. S7B). However, when 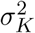 increases enough that *σ*_inv_ ∼ 1/ℛ_0_, the distribution of fitness effects will be somewhat distorted from the Gaussian prediction (Fig. S7C). This is because resource availabilities *h*_*i*_ are non-negative and can only be approximated by a Gaussian when Std(*h*_*i*_) ≪ ⟨*h*_*i*_⟩ = 1.

#### 4.2 Variation in Number of Metabolized Resources

In the binary resource model so far, we have assumed that each resource has an independent probabilityℛ _0_/ℛ of being used by each species. This assumption results in each species using a binomially distributed number of resources, which is tightly peaked when ℛ_0_ ≫ 1. However, there is no reason to assume that coexisting species cannot differ in their levels of specialism, such that some use many resources and some few. First, it is useful to consider what happens if the parent species uses ℛ_*P*_ resources on average, rather than the ℛ_0_ resources that most species in the community use. This extension helps us disentangle what terms in our expressions depend on the typical metabolic overlap between members of the community (encoded by ℛ_0_), and what depends on the specific number of resources used by the parent and mutant species. It can also help determine whether generalist or specialist species are more likely to diversify in a given community.

The more resources a given organism uses, the more its growth rate and overall fitness are averaged out between different independent variables, effectively reducing the variance in its abundance as well as the invasion fitness of any particular mutation. So, if ℛ_*P*_ is larger, the parent species is less likely to have particularly high abundance, but the invasion fitness of a strategy mutation it obtains is likely to be smaller in magnitude. The first effect should lower the coexistence probability, while the second should raise it. Performing the calculation up to our expression for the joint probability of Δ_*P*_ and Δ_*M*_ gives us

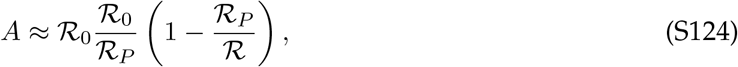

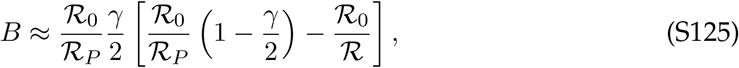

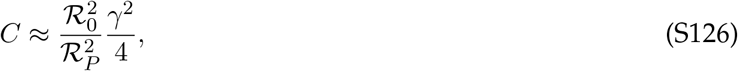

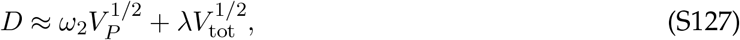

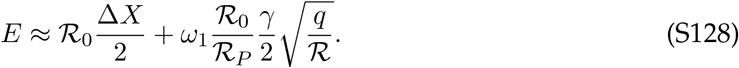

Here, we have defined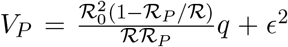, reflecting the different variation in overall fitness between the parent and other species in the community. We have once again defined *γ* such that *γ* ≈ 1 for a knock-out mutation.

Since most of the community is unaffected by changing ℛ_*P*_, we use the same saddle-point results we calculated before. Thus, the value of *σ*_inv_ is unchanged, although the invasion fitness of the mutant has a prefactor of ℛ_0_/ℛ _*P*_ to account for the altered number of resources. The inequality we obtain that predicts mutant-parent coexistence (for Δ*X* = 0) is now given by

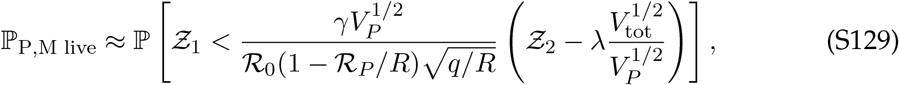

where 𝒵_1_ is once again proportional to the invasion fitness of the mutant, and the 𝒵_2_-dependent factor on the RHS is proportional to the abundance of the parent strain. The calculations otherwise proceed as before, yielding the results shown in Figure S9A. For these parameter values, it appears that mutant-parent coexistence is more likely for mutations on a background strain which metabolizes many resources, since these mutations tend to have small invasion fitness.

Next, we consider what happens when the number of resources ℛ_0_ utilized by *every* species is more broadly distributed, to allow coexistence of generalists and specialists. Specifically, we consider ℛ_0_ ∼ Uniform(20, 130) with ℛ = 150. The actual number of resources used by surviving species will not be uniformly distributed because specialists are more likely to have particularly “lucky” resource strategies; similarly, the typical number of resources used by species which produce a successful knock-out mutant will also be different (Fig. S9C). Rather than analytically calculating all of these distributions for this somewhat artificial example, we can use the observed distribution of ℛ_0_ and ℛ_*P*_ as empirical inputs to our analytic predictions of fitness effects and mutant-parent coexistence (in particular, plugging in the mean value of ℛ_0_, and integrating over the probability distribution of ℛ_*P*_). Doing so yields reasonable predictions for the mutant-parent coexistence probability and distribution of fitness effects (Fig. S9B,D), confirming that our results do not critically depend on each member of the community utilizing a similar number of resources.

#### 4.3 Metabolic Trade-offs in Sampling

In our model, we have posited a soft metabolic trade-off where each sampled organism *µ* has an identically distributed “uptake budget” 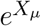 which must be divided between the resources it consumes; thus, organisms which consume more resources are generally less effective at consuming any individual one. Other work has considered different forms of such trade-offs (37, 44) or none at all, which can alter the number of species that coexist in the community (29). For example, Refs. (36) and (37) also consider a soft metabolic trade-off, but where the cost of consuming additional resources is associated with a larger average death rate rather than with taking a portion of a fixed overall budget. While these models describe very similar final communities, the main difference is that the death rate trade-off model predicts that as sampling depth 𝒮/*𝒮* ^*^ increases, the fitness effect of using an additional resource becomes slightly deleterious on average, such that the mean invasion fitness of knock-out mutations is small but positive (Fig. S8A). Accounting for this small fitness effect through an effective change in pure fitness Δ*X* in knock-out and knock-in mutants brings the mutant-parent coexistence probabilities in agreement between the models (Fig. S8B).

Another alternative is to eliminate metabolic trade-offs in sampling entirely (29). In terms of the model we have discussed so far, a lack of trade-offs effectively grants organisms which use more resources a larger budget to divide among those resources. This change causes organisms which use more resources to be overrepresented in the population due to their higher effective fitness, and lowers the niche saturation 𝒮^*^/*ℛ* of resultant communities. However, when these empirical changes in effective ℛ_0_ and 𝒮^*^/*ℛ*are accounted for, the distribution of fitness effects and mutant-parent coexistence probability are well-described by the original model with trade-offs (Fig. S8C-D). It is important to distinguish metabolic trade-offs in *sampled* species from trade-offs in mutations, which can be captured by changing the value of Δ*X* as described in the main text. These trade-offs may be quite different over short evolutionary timescales: for example, first-step mutations might introduce metabolic inefficiencies which are reduced by subsequent mutations.

#### 4.4 Continuous Resource Usage

We have generally considered a binary model of resource use, where a natural parameter to describe metabolic overlap is the number of resources ℛ_0_ used by a typical species. A simple continuous analogue to binary resource usage is drawing resource strategies from a Dirichlet distribution, 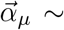 Dirichlet (ℛ_0_/ℛ, 1). Typical resource consumption vectors sampled from this distribution are qualitatively similar to binary ones; in particular, they will have ∼ ℛ_0_ entries which are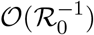, with the remaining entries much smaller. As an analogue to knock-out mutations, we chose a random entry of the parent organism’s strategy vector with value at least *γ*/ℛ _0_, and reduced it by *γ*/ℛ _0_, where *γ* = 1 represents a full knock-out. We also considered “global-effect” mutations which slightly perturb each element of the resource use vector. Specifically, a global strategy mutation 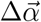 was constructed by multiplying each element of 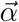 by an independent Gaussian random variable with mean 0 and variance 1/ℛ_0_, then by rescaling such that ∑_*i*_ Δ*α*_*i*_ = 0 and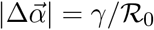.

Using resource consumption vectors drawn from a continuous distribution in this manner left the quantitative properties of our communities relatively unchanged (Fig. S3A-B, Fig. S5).

As a more extreme example of continuous resource usage, we can consider a community where all resources are significantly utilized by all species, but at slightly different levels.

For example, we can draw 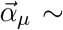 Dirichlet(*B*, 1) for *B* ≫ 1. In this case, the distribution of resource consumption strategies will approach a Gaussian,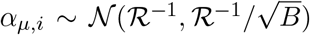; similar sampling schemes have been analyzed before, as in (29). With these Gaussian resource consumption strategies, the number of resources ℛ_0_ utilized by a typical species no longer has intuitive meaning. However, we can generalize our conclusions from binary resource usage by defining an effective ℛ_0_ such that the binary model and the Gaussian model predict the same level of variance 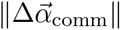 in resource consumption strategies:

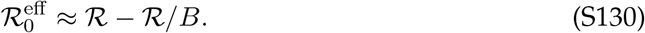

Thus, a model where organisms use all resources at slightly different levels is equivalent to a binary resource usage model where organisms use *almost* all resources, in terms of the metabolic overlap between species. We can define mutations in this framework by perturbing the parent’s resource usage 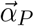 toward a different, i.i.d. resource usage profile 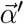:

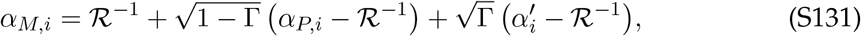

where Γ ≤ 1 is a mutation effect size parameter describing how much of the initial resource use profile 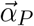 is retained.

What should the parent-mutant coexistence probability be for such a mutation? Plugging in the 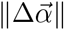 and 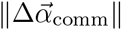 for this assembly process into Eq. (9), we find

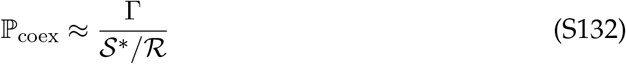

for Γ ≪ 1. Indeed, we find that the mutant-parent coexistence probability for this community is well-predicted by naively using the binary resource predictions with the value of ℛ_0_ defined in Eq. (S130), and has the expected qualitative dependence on Γ and *B* (Fig. S3C).

#### 4.5 Specialist Community Assembly

As discussed in SI Section 3.5, the replica-theoretic approach involves a “large-ecosystem” approximation which is only valid when 𝒮^*^ℛ_0_ *>*ℛ, such that each resource is consumed by many species. Species coexisting in such ecosystems tend to not have large differences in their uptake budgets *X*_*µ*_, although these differences can still be ecologically relevant. Here, we consider a simple “specialist assembly” procedure, as a representative example that violates this condition. Imagine that an initial community contains 𝒮^*^ species, each of which use exactly one resource with no overlap. These species will trivially coexist, at abundances which match the supply of their private resource, *n*_*µ*_ = *K*_*µ*_. Then, suppose that one organism *P* acquires a partial knock-in mutation which enables it to use another resource *ν*. We label the mutant *M*, so that

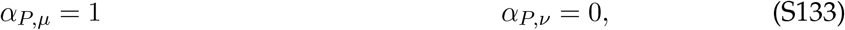

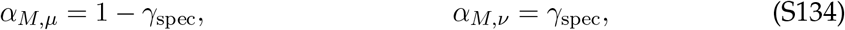

where *γ*_spec_ describes the effect size of the mutation. While this ecosystem is clearly a caricature, it is a useful example far from the regime where our results should be applicable with which to compare our qualitative results.

What is the impact of the mutation? If no other organism utilizes resource *ν*, then the mutant will trivially always survive and coexist with its parent. However, if another organism *O* is already utilizing resource *ν*, then the mutant will survive if its fitness is greater, *X*_*M*_ *> X*_*O*_. If the mutant survives, it will either replace its parent strain, or coexist with it while replacing *O*. Specifically, mutant-parent coexistence occurs if the following condition is met:

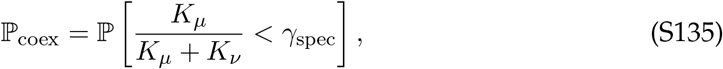

as well as the trivial case where no other organism utilizes *ν*. Although the condition for coexistence is different from the more complex assembly procedure we discuss elsewhere, we still find that it is nonzero even in fully saturated communities (Fig. S3D). Thus, while our results do not directly extend to these communities, their existence does not invalidate our main conclusions. Furthermore, we expect that the sparser interaction networks of these communities makes them more amenable to previous analysis focusing on small numbers of species and resources (30).

### 5 Simulations and Numerics

#### 5.1 Community Simulations

Throughout this work, we have performed numerical simulations of community assembly in order to verify and extend our analytical results. These simulations were performed by randomly sampling species and solving the optimization problem corresponding to ecological equilibrium (described in SI Section 1.5), rather than explicitly solving the abundance dynamics: the final outputs are the resource availabilities *h*_*i*_ and the surpluses Δ_*µ*_. The code for these simulations is based on that in Ref. (37).

Much of our analysis relies on identifying whether individual species are present in the population or extinct. Mathematically, these cases correspond to whether Δ_*µ*_ is zero or negative. Numerically, however, Δ_*µ*_ is never precisely zero, necessitating the definition of an “extinction threshold.” Any species with a surplus Δ_*µ*_ below the extinction threshold, a small negative number, is classified as extinct. We typically defined the extinction threshold as − 10^−3^ · Std(*h*_*i*_)/*ℛ* _0_, several orders of magnitude smaller than the typical fitness effect of a knock-in or knock-out mutation.

In Figure S10, we demonstrate that this is a reasonable choice of threshold. In order to be numerically sensible, there should be a clear separation between typical values of Δ_*µ*_ for surviving and extinct species, and the threshold should be well within this gap. We verified this by plotting the number of putative surviving species as a function of the extinction threshold. In the neighborhood of the threshold we have chosen, the number of putative survivors is independent of the threshold, suggesting that there is a clear separation between alive and extinct species and that our choice of threshold should not heavily impact our results. We also analyzed the values Δ_*µ*_ for each strain before and after mutant invasion in order to ensure that we were reliably identifying species that go extinct. A scatterplot of initial vs. final Δ_*µ*_ values shows that almost all species are clearly separated into a “surviving” cluster and an “extinct” cluster, again suggesting that our results are not due to numerical artifacts.

#### 5.2 Theory Numerics

While the main text and remainder of the SI describe the novel theory key to our work, some theory curves shown in figures involve minor post-processing steps or slightly more precise calculations than those quoted in the text. For convenience, we summarize these steps (which are also performed in our code supplement) below.

##### Distribution of fitness effects (e.g. Fig. 2C-D)

Our theoretical results predict that the distribution of fitness effects (DFE) follows a Gaussian distribution with mean zero and standard deviation *σ*_inv_, which is calculated by plugging our parameters into the equations in SI Section 3.2. The mean of this distribution is taken as a variable Δ*X* (sometimes assumed to be zero).

##### Coexistence probability (e.g. Fig. 3)

We calculate ℙ_coex_ by analytically integrating Eq. (S107) over the probability distributions of *s*_inv_ and *Ƶ*_2_. Both distributions are truncated Gaussians multiplied by a linear term: *p*(*s*) ∼ *sρ*(*s*)*θ*(*s*), where *ρ*(*s*) is the distribution of fitness effects above and *θ*(*s*) is the Heaviside function. Z_2_ is related to the parent relative abundance via Eq. (S101), and thus takes on a similar distribution: 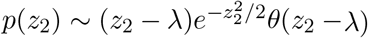. In the inset of Fig. 3D, ℙ_coex_ is conditioned on *s*_inv_, so it is not integrated over. In Fig. S6, we condition on both *s*_inv_ and _2_ (via *f*_*P*_), so Eq. (S107) determines a line in the *s*_inv_-*f*_*P*_ plane above which mutant-parent coexistence is permitted.

##### Relative abundance (e.g. Fig. 4D)

Relative abundance *f*_*µ*_ is calculated from simulation output by identifying surviving species using the resource surplus Δ_*µ*_ (as described in SI Section 5.1) and then using a non-negative least-squares solver to invert the relationship 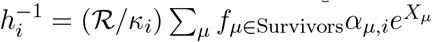.

##### Simluating model extensions (e.g. Fig. S3C)

For some extensions of the model, the theoretical expectation of 𝒮^*^/*ℛ* was not explicitly calculated. Rather than choosing parameters to target a specific value of 𝒮^*^/*ℛ*, simulations were run with a fixed value of Std(*X*_*µ*_) and varying log-spaced 𝒮/ *ℛ* to cover a range of 𝒮^*^/*ℛ*. To make smooth theoretical predictions from this data, the empirical values of *λ*(𝒮^*^/*ℛ*) were linearly interpolated over the sampled data. In Fig. S8C-D, the sampled R_0_ differed from the R_0_ among survivors due to the systematic advantage in using more resources; as such, the empirical average _0_ among surviving species was used as input to the theory.

*Figure 2B*: We numerically solve the system of equations in SI Section 3.2 for *λ* and find 𝒮^*^/*ℛ* by inverting Eq. (S88).

*Figure 2E*: We calculate the correlation between the expressions in Equations (S120) and (S121) in terms of *σ*_inv_.

*Figure S1*: The set of equations in Eq. (S1a) were numerically solved using a standard MATLAB differential equation solver, with random initial conditions corresponding to low species abundances and resource concentrations at their steady-state in the absence of species.

*Figure S8A-B*: An effective Δ*X* = *m*/*ℛ* is determined by Eq. (S89) with *χ* = 0.

*Figure S9A*: In the left panel, ℙ_coex_ is determined by integrating Eq. (S129) over its random variables rather than Eq. (S107). Z_1_ is a standard Gaussian truncated Jto take positive val-ues, while Z_2_ is a standard Gaussian truncated to take values above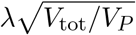.

*Figure S9B-D*: In addition to the changes discussed in “Simulating model extensions” above, the mean values of ℛ_0_ and ℛ_*P*_ among surviving species were linearly interpolated over the sampling regions of each condition to produce the theory curves in panel B. While the mean value of ℛ_0_ was used for the DFE in panel D, the entire empirical distribution of ℛ_*P*_ (i.e., the number of resources used by species which were randomly drawn to produce mutants across all simulation trials) was numerically integrated over to account for the tail effects.

